# RNA Dynamics Regulate Transcriptional Condensate Vivacity to Drive Gene Coordination

**DOI:** 10.1101/2025.06.06.658161

**Authors:** Koichi Ogami, Ming M. Zheng, Seiko Yoshino, Yohei Sugimoto, Mayu Seida, Toshiki Mori, Ryo Yoshizawa, Koh Onimaru, Mitsutaka Kadota, Shigehiro Kuraku, Kenichi G. N. Suzuki, Ibrahim I. Cissé, Phillip A. Sharp, Hiroshi I. Suzuki

**Author notes:** These authors contributed equally to the work (Koichi Ogami, Ming M. Zheng). To whom correspondence should be addressed., Division of Molecular Oncology, Center for Neurological Diseases and Cancer, Nagoya University Graduate School of Medicine, 65 Tsurumai-cho, Showa-ku, Nagoya 466-8550.

## Abstract

Transcriptional condensates (TCs), enriched with Mediator, orchestrate super-enhancer (SE)-driven gene expression critical for cell identity. However, how their dynamic physical properties shape transcriptional activities remains unclear. Here, we reveal a previously unknown regulatory axis wherein dynamic features of enhancer RNA (eRNA), transcribed but rapidly degraded by the RNA exosome, maintain optimal TC fluidity and stability. Depletion of RNA exosome disrupts TC integrity, altering Mediator and Pol II colocalization, in embryonic stem cells. This TC perturbation attenuates proper transcriptional regulator loading and pause-release regulation, enhances transcriptional noise, and diminishes coordinated eRNA-mRNA expression for SE-associated genes. Multi-omics analyses, live-cell imaging, and computational modeling collectively demonstrate that RNA turnover modulates TC vivacity, facilitating widespread chromatin contacts between SE-containing active A compartments and coordinated transcriptional bursts across SE-associated genes. Our findings establish RNA-dependent condensate dynamics as an essential quality-control mechanism that fine-tunes global transcriptional coordination, maintaining cellular homeostasis and cell identity.

**Highlights:**

- RNA exosome is required for the homeostasis of transcriptional condensates (TCs)
- RNA synthesis and decay optimize the fluidity and dynamic properties of TCs
- TCs maintain the transcriptional consistency of super-enhancer (SE) genes
- TCs support contacts of SE-containing A compartments for coordinated transcription

## INTRODUCTION

Proper operation of distinct gene expression programs in diverse cell types is crucial in multicellular organisms.^1^ Such cell type- or tissue-specific gene expression programs are orchestrated by complex interplays between cell type-specific transcription factors (TFs), including so-called master TFs, enhancer activities, and epigenetic regulation. These programs are also modulated by hierarchical chromosome structures, which include A/B compartments, topologically associating domains (TADs), and enhancer-promoter loops^2^. Further, transcription of individual gene is regulated through multiple checkpoints of RNA polymerase II (Pol II): initiation, pause, elongation, and termination, and exhibits bursting features with varying degrees.^3–5^

Super-enhancers (SEs) are a class of regulatory regions characterized by clusters of enhancers with exceptionally high occupation of master TFs and coactivators, including Mediator and BRD4, and high-level histone H3K27ac modification.^6^ Several hundreds of SEs can be identified by using ChIP-Seq datasets for master TFs, Mediator, and/or H3K27ac. SEs activate genes important for functionality of the corresponding cell type, which include master TFs and miRNAs, forming a feedback loop and ensuring cell identity.^6–8^ Expanded from studies in mouse embryonic stem cells (mESCs), SEs have been identified in various types of cells, including cancer cells, and linked to the pathogenic mechanisms of various diseases, including cancer.^7,9,10^ SEs include multiple functional constituents,^11–14^ involve complicated genomic interactions among multiple SE constituents and promoters,^15–18^ and are often associated with the formation of TAD triplets, suggesting high-order structures beyond simple enhancer-promoter loops.^19,20^

Membraneless subcellular organelles, termed biomolecular condensates, such as the nucleoli, germ granules, and the Cajal body, are formed by phase transitions and organize biochemical processes in cells.^21,22^ Recent evidence has revealed similar microscopic condensates concentrating large numbers of TFs, cofactors, and Pol II, termed transcriptional condensates (TCs), are associated with the transcription of SE-associated genes.^23,24^ Mediator, BRD4, Pol II, master TFs, and other transcriptional regulators possess intrinsically disordered regions (IDRs),^23,25,26^ which promote multivalent interactions critical for condensate formation. Mediator and Pol II form large stable (> 300 nm) and small transient bodies (∼100 nm) in the nucleus of mESCs, with dynamic properties of condensates.^24^ FISH and live-cell imaging analyses have shown that SE-associated genes are incorporated into TCs.^23–25^ More recently, live-cell imaging focusing on spatial distribution of Sox2 as a representative SE-controlled gene in mESCs revealed that diffraction-sized TCs (∼400-500 nm) exist in the proximity of Sox2 gene, and transcriptional bursting of Sox2 gene occurs when the nearest TC moves within < 1 µm from Sox2 gene.^27^ These findings suggest TCs are the critical molecular machinery for SE-mediated transcription.

The properties of condensates provide explanation of certain features of enhancers: the sensitivity of SEs to perturbation, the transcriptional bursting patterns of enhancers, and the ability of an enhancer to activate multiple genes, as observed in *Drosophila*.^28–30^ However, it remains largely unclear how qualities and properties of TCs are dynamically regulated and how such regulation is linked to functional aspects of transcription, including the relationships with genome structures, stepwise regulation of Pol II, and transcriptional coordination of different genes. Growing evidence has revealed that RNA molecules are key determinants in the formation of many cellular condensates, including nucleoli and paraspeckles.^31–36^ Importantly, TCs are proposed to show RNA-driven reentrant phase transition behavior: low-level RNAs transcribed from enhancers (enhancer RNA, eRNA) promote an initial condensate formation, but excessive RNAs produced by transcriptional burst form a negative feedback mechanism.^37,38^ This RNA feedback model can be explained by electrostatic balance between RNAs and proteins.^37^ As potential RNA sources of a facilitator of initial TC formation, enhancers and promoters are transcribed into bidirectional eRNAs and upstream antisense RNAs (uaRNAs), respectively. eRNAs and uaRNAs have several important features: generally short (< 1 kb), chromatin-associated, and unstable (eRNA t_1/2_ ∼ a few minutes; uaRNA t_1/2_ ∼ 1-10 min) due to activities of the RNA exosome, a 3′-5′ exonuclease complex.^39–42^ These observations suggest that, in addition to RNA synthesis, rapid destabilization of eRNAs by RNA exosome could be important in condensate homeostasis.

By combining physics-based modeling and experimental analysis, we here propose that RNA dynamics regulate the material properties and dynamic functions of transcriptional condensates. We first explored the impacts of short-lived RNA decay on Mediator-enriched TCs by utilizing RNA exosome conditional knockout (CKO) mESCs.^43^ Consistent with our hypothesis, depletion of Exosc3, a core component of RNA exosome, led to disappearance of large TCs (> 0.3 µm^2^) that overlap with Pol II clusters and an increase of smaller condensates. We validated the robustness of our RNA exosome CKO cell model system, allowing systematical analysis of the impacts of TC dynamics on SE functions. With computational simulations, live-cell imaging, and multi-omics analyses, we found that RNA synthesis, decay, and mobility contribute to not only TC formation but also enhancement of their fluidity and dynamic properties, which we collectively call “TC vivacity”. Importantly, we found that TC vivacity is important for three aspects of coordinated transcription of SE-associated genes: suppression of transcription noise, eRNA-mRNA co-expression, and gene-gene co-expression associated with pervasive intrachromosomal interactions. Together, our findings illustrate novel roles of TCs as a quality control system of coordinated transcription.

## RESULTS

### RNA exosome as a key player in RNA feedback model

The current RNA feedback model proposes that low levels of short RNAs (e.g., eRNAs) enhance TC formation, while high levels of longer RNAs (pre-mRNAs) promote condensate dissolution, driving transcription burst cycles (Figure 1A).^37^ Given that MED1-IDR undergoes reentrant phase transition in response to increasing short RNA levels,^37^ we hypothesized that eRNA stabilization by RNA exosome loss could suppress TC stability (Figure 1A). To test this, we engineered previously established Exosc3 CKO mESCs^43^ using CRISPR-Cas9 to tag endogenous MED19 with a Halo-tag.^24^ MED19 and Exosc3 are subunits of the Mediator and RNA exosome complex, respectively. While RNA exosome depletion gradually causes differentiation and cell death in FBS/LIF culture conditions in our report,^43^ notably, Exosc3 depletion caused neither morphological changes nor differentiation phenotypes in 2i culture (Figure S1A). This was further confirmed by subsequent single-cell transcriptome analysis (see later results in Figure 5). 2i conditions have been used in previous TC studies,^23,24^ and are suitable for microscopic observations of TCs. Consistent with our hypothesis, super-resolution imaging with photo-activation localization microscopy (PALM) disclosed that Mediator condensates shrink upon RNA exosome depletion (Figures 1B and S1B).

**Figure 1.**
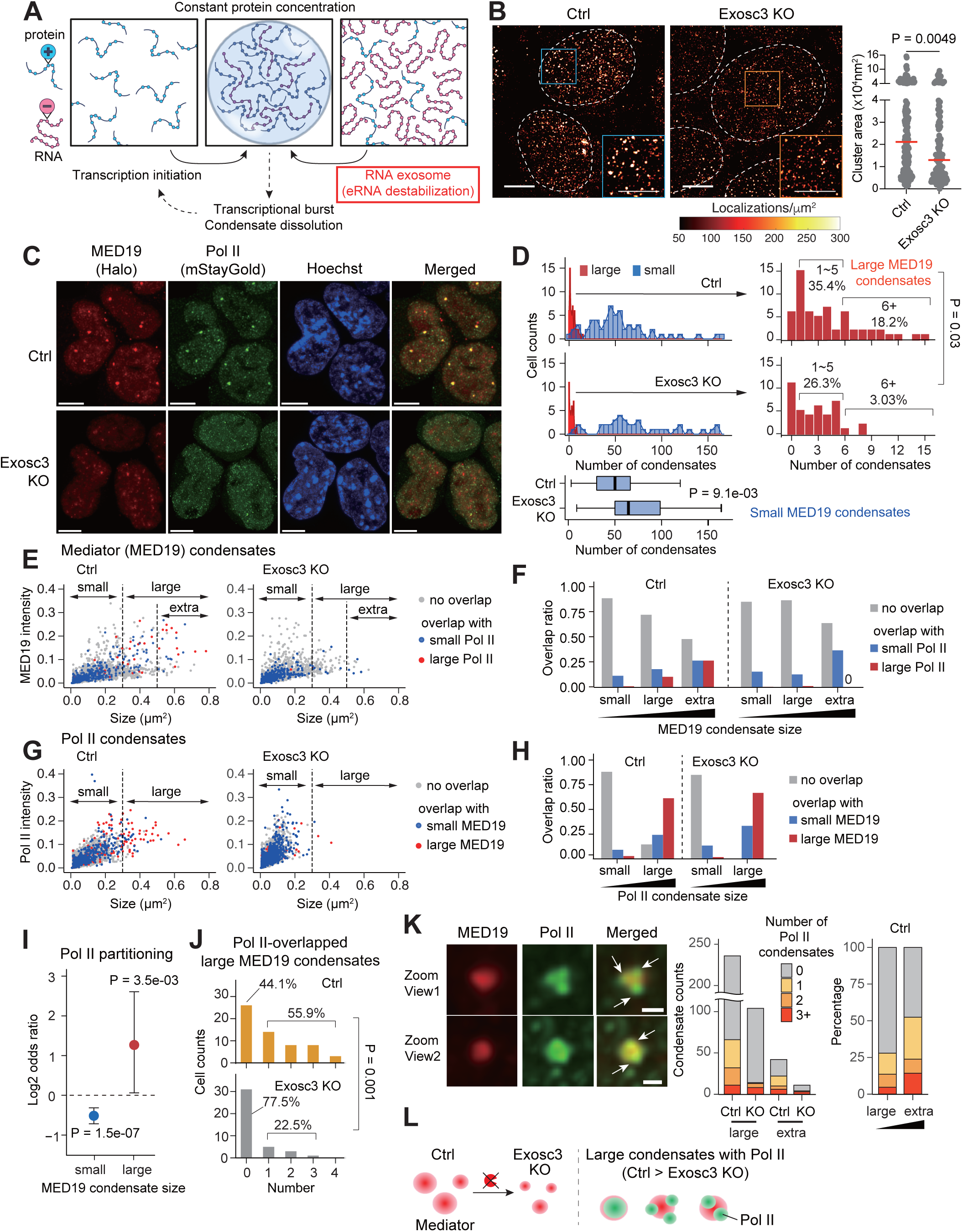
Expansion of RNA feedback model: loss of RNA exosome disrupts transcriptional condensate homeostasis. (A) A schematic illustrating reentrant phase transition driven by the balance between RNA and protein, derived from *Henninger et al.*^37^ Transcription initiation promotes formation of charge-neutralized condensates at low to intermediate RNA:protein ratios. Transcriptional bursts lead to condensate dissolution, forming RNA feedback cycles. However, at super-stoichiometric RNA concentration, the condensate formation is disrupted. The RNA exosome is hypothesized to enable condensate formation by preventing excess eRNA accumulation. (B) PALM super-resolution imaging of endogenous Mediator labeled with (PA)JF549-HaloTag in Exosc3 CKO mESCs. The right jitter plots show the area of each cluster. Scale bars, 4 µm; P value, Wilcoxon rank sum test. (C) Confocal microscopic images showing the localization of JF646-Meditor (MED19) and mSG-Pol II under Ctrl and Exosc3 KO conditions. Scale bars, 5 µm. (D) Histograms showing the number of small (< 0.3 µm^2^) and large (≥ 0.3 µm^2^) MED19 condensates per nucleus. Bottom boxplots and right bar graphs represent distribution of small and large condensate counts per nucleus, respectively. P value, Wilcoxon rank sum test (box plots) and Fisher’s exact test (bar plots). (E) Scatter plots showing the relationships between MED19 condensate signal intensity and size (small (< 0.3 µm^2^), large (≥ 0.3 µm^2^), and extra-large (≥ 0.5 µm^2^)). Blue and red dots indicate overlap with small (< 0.3 µm^2^) and large (≥ 0.3 µm^2^) Pol II foci, respectively. (F) Bar plots summarizing the overlaps in (E). (G, H) The relationships of Pol II foci signal intensity and size and MED19 overlap, as in panels (E) and (F). (I) Forest plots showing the log-scaled odds ratio of Pol II partitioning in small and large MED19 condensates. Bars, 95% confidence intervals; P value, Fisher’s exact test. (J) Bar graphs showing the number of large MED19 condensates co-partitioned with Pol II. P value, Fisher’s exact test. (K) Zoomed-in view of Mediator and Pol II overlap condensates. Arrows mark the positions of Pol II condensates (left). Bar plots summarize the number of Pol II condensates per Mediator condensate (right). Scale bars, 500 nm. (L) A schematic depicting the formation of MED19 condensates co-partitioned with Pol II condensates. Exosome loss reduces the formation of large MED19-Pol II condensates. See also Figure S1.

### Loss of RNA exosome disrupts large Mediator-Pol II condensates

Given that Mediator and Pol II form large stable condensates associated with chromatin,^24^ we next analyzed colocalization of Mediator and Pol II by tagging endogenous POLR2A (RPB1) with mStayGold (mSG)^44^ in Exosc3 CKO/Halo-MED19 cells. Using confocal microscopy, we found that Exosc3 depletion leads to the loss of large Mediator condensates associated with Pol II and an increase in smaller condensates (Figure 1C). We further performed cell image analysis (Figure S1C) to quantitate their characteristics and colocalization (Figures 1D-J and S1D-G). First, for both Mediator and Pol II, a decrease in large condensates (> 0.3 µm^2^), an increase in small condensates, and an overall decrease in size and intensity were observed upon Exosc3 depletion (Figures 1D and S1D-F). In size-dependent colocalization analysis under control conditions, as Mediator condensates enlarge, they frequently associate with Pol II, consistent with the previous report (Figures 1E-F).^24^ Importantly, following Exosc3 depletion, Pol II preferentially dissociates from large Mediator condensates (Figures 1E-F). These findings are supported by Pol II partitioning analysis (Figure 1I) and separate quantitation of large Mediator-Pol II condensates (Figures 1J and S1G).

While RNA exosome depletion disrupts most of large Pol II condensates (Figure 1G), we observed that remaining large Pol II condensates are associated with Mediator condensates at a similar frequency (Figure 1H). Closer inspection of large Mediator-Pol II condensates unveiled that these condensates accommodate multiple Pol II clusters, with the number correlating with the condensate size and the presence of Exosc3 (Figure 1K). Consistent with the maintenance of pluripotent states, alterations in MED19 condensate formation and Pol II co-partitioning were reversibly rescued by re-induction of Exosc3 expression (Figure S1H). In addition, growth suppression by Exosc3 depletion in 2i media was reversed by adding back Exosc3 (Figure S1I). Collectively, these results indicate that RNA destabilization by RNA exosome plays key roles in maintaining TC stability, particularly large ones (Figure 1L).

### RNA tunes the fluidity and physical properties of TCs

In contrast to RNA exosome depletion, inhibition of RNA elongation reportedly enhances TC size (Figure S2A).^37^ Given the reentrant phase behavior of RNA-protein mixtures, the steady-state TCs in cells are presumably optimized by the balance between RNA synthesis and RNA decay for normal TC function and gene expression (Figures 2A and S2A). This assumption is also based on the function of RNA exosome preventing deleterious R-loop structures for proper enhancer activities.^39^ Moreover, TC size (associated with TC stability) only manifests one aspect of molecular interactions which underpin many other condensate properties. Therefore, we next investigated how RNA synthesis, decay, and mobility, three key features of enhancer transcription dynamics, affect the quality of TCs from different aspects. To do so, we adapted a physics-based model to evaluate various physical properties of TCs, including intensity, sharpness, incorporation rate, intra-condensate molecule exchange, and chromatin accessibility (Figures 2B and S2) under the dynamic perturbations in RNA synthesis, degradation, and mobility (Figure 2C) (STAR Methods). The intensity changes in simulation correspond to TC size changes in the experiments. Simulation results based on a general Landau approach, an extended model from the RNA feedback model study,^37^ recapitulated increased and decreased TC intensity upon transcription inhibition and RNA exosome depletion, respectively (Figure S2B). Further, the simulation predicteded that both transcription inhibition and RNA exosome depletion attenuate intra-condensate molecule exchange and incorporation rate, indicative of reduced TC fluidity (Figures 2D and S2C-F, and Video S1) (STAR Methods).

**Figure 2.**
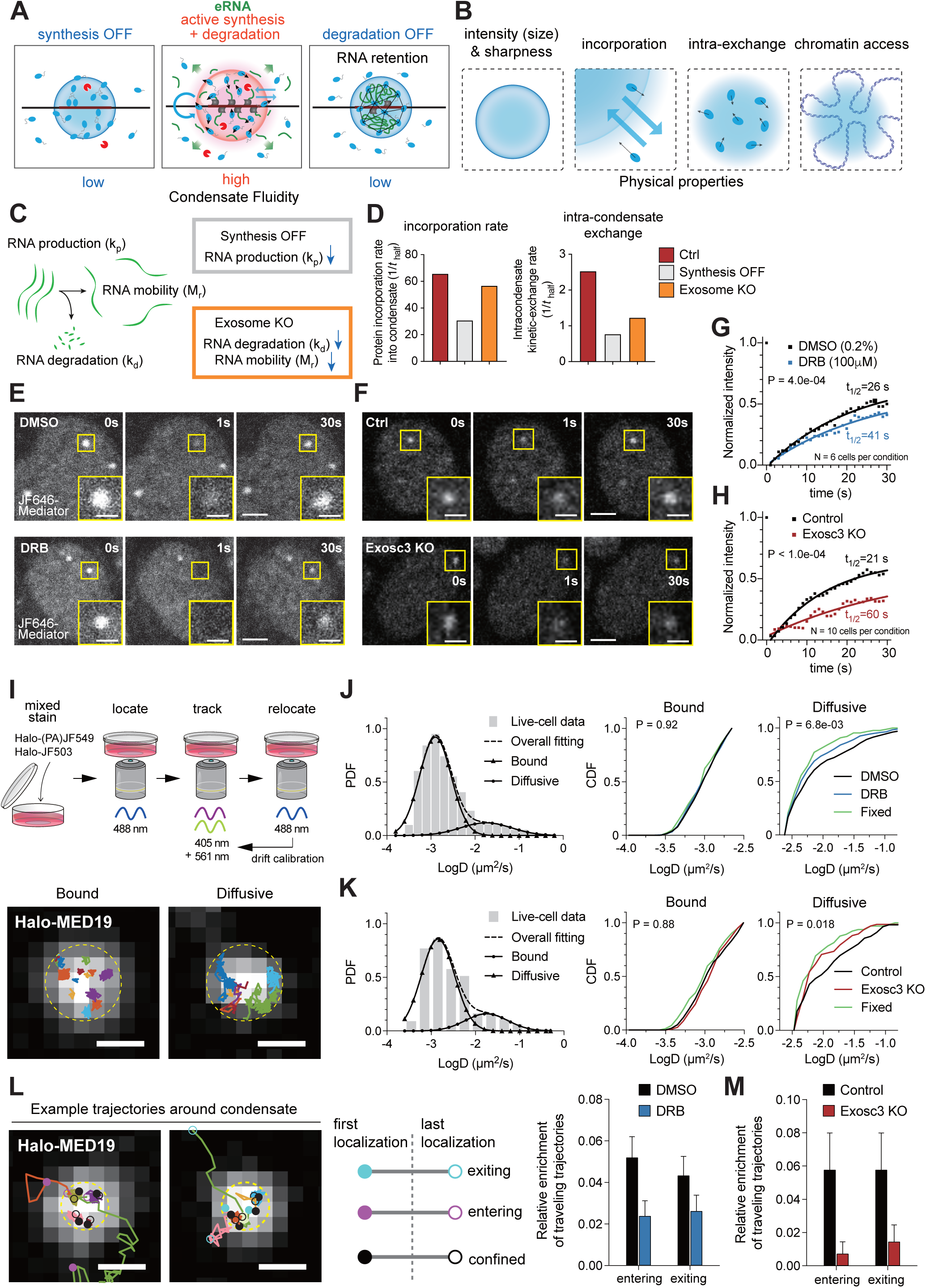
RNA synthesis and destabilization fluidize transcriptional condensates and modulate their physical properties. (A) A scheme depicting the effects of inhibition of RNA transcription and destabilization on condensate fluidity. (B) Hallmark properties of transcriptional condensates, which is analyzed by phase-field simulations. (C) The model framework of RNA dynamics. The schematic derived from *Henninger et al.*^37^ k_p_, RNA production rate; k_d_, RNA degradation rate; M_r_, RNA mobility. (D) Simulation results showing the effects of inhibition of RNA transcription (synthesis OFF) and Exosome KO on the condensate incorporation rate and intra-condensate exchange rate. Details are shown in Figure S2. (E, F) FRAP analysis of Mediator condensates upon 100 µM DRB treatment (20 min, vs DMSO) (E) and Exosc3 depletion (F). Images show nuclei before (left, 0 sec), immediately after (middle, 1 sec), and after bleaching (right, 30 sec). The boxes indicate the bleached positions, and enlarged views of the bleached areas are shown. (G, H) Normalized recovery curves for Mediator in DRB treatment (D, n = 6, 20 min treatment) and Exosc3 depletion (E, n = 10), respectively. The estimated half-recovery times (t_1/2_) are also shown. P value, two-tailed *t*-tests with Welch’s correction. (I) Scheme depicting single-molecule tracking of Halo-MED19 using mixed staining with Halo-(PA)JF549 and Halo-JF503 (top). Representative trajectories in bound and diffusive states with bulk staining of MED19 condensate as background (bottom). Dashed lines, condensate boundaries; Scale bars, 500 nm. (J) Histogram of diffusion rates (log D) and probability density function (PDF) curves for overall, bound, and diffusive trajectories (left). CDF plots of diffusion rates for bound and diffusive trajectories in live cells treated with DMSO or DRB (100 µM, 20 min), as well as in paraformaldehyde-fixed cells, are shown (right). P value, two-tailed *t*-tests with Welch’s correction. (K) Single-molecule tracking analysis in live control, Exosc3-depleted, and fixed cells, as in panel (J). P value, two-tailed *t*-tests with Welch’s correction. (L) Traveling trajectory analysis. Trajectories were categorized into three groups (left): exiting, entering, and confined. The bar plot shows the relative enrichment of entering and exiting trajectories in cells treated with DMSO or DRB (100 µM, 20 min) (right). Dashed lines, condensate boundaries; Scale bars, 500 nm. (M) Bar plot showing the relative enrichment of entering and exiting trajectories in control and Exosc3 KO cells. Error bars: SEM. See also Figures S2 and S3.

To experimentally test these predictions, we first measured the rate of fluorescence recovery after photobleaching (FRAP) for Halo-MED19-labeled TCs (Figures 2E-H). Supporting the model’s predictions, recovery from photobleaching was delayed by 5,6-dichlorobenzimidazole riboside (DRB)-treatment (Figure 2E), which disrupted transcription elongation, and by Exosc3 depletion (Figure 2F), with 1.6-fold (DRB vs DMSO) and 2.9-fold (Exosc3 KO vs control) prolongation in recovery half-times (Figures 2G-H). We next performed live-cell single-molecule tracking of Halo-MED19 under mixed staining with two different probes, with JF503 for visualizing the bulk distribution and (PA)JF549 for molecule tracking (Figure 2I). Diffusion rate profiles suggested the presence of stationary (bound) and mobile (diffusive) populations (Figures 2J-K). Both populations of MED19 can associate with the Nanog transcription site (Figures S3A-B), as previously suggested for transient interaction between gene locus and a condensate.^24,27^ Consistent with the predictions, the diffusion rate of diffusive MED19 molecules decreased under DRB treatment (Figure 2J) and further decreased with Exosc3 depletion (Figure 2K), approaching the pseudo diffusivity due to localization uncertainty (see the paraformaldehyde-fixed condition). Furthermore, detailed trajectory analyses of MED19 molecules entering and exiting condensates revealed that both events were less frequent (Figures 2L-M), and the dwell time of MED19 molecules in condensates increased in DRB-treated and Exosc3-depleted cells (Figures S3C-E). Similarly, single-molecule tracking of Pol II with respect to MED19 condensates showed that Pol II entering-in events into MED19 condensates were less frequent in Exosc3-depleted cells (Figures S3F-G, and Video S2).

### RNA modulates the sharpness and chromatin accessibility of TCs

Our single-molecule trajectory analysis suggests a steep separation of the protein diffusion dynamics at the condensate interface in DRB-treated and Exosc3-depleted cells, which presumably echoes with increased boundary sharpness and decreased chromatin accessibility. Consistently, our models predicted both effects (Figures 3A-C and S3H). (STAR Methods). TC and chromatin were predicted not to be mixed well upon Exosome depletion (Figure 3C). We then experimentally validated the predictions by imaging analyses (Figures 3D-J). Measurement of signal intensity profiles along the condensates allowed us to confirm that boundary sharpness of MED19 condensates was indeed significantly increased upon DRB treatment and Exosc3 depletion (Figures 3D-F). Chromatin accessibility was assessed by overlaps of the condensates with H2B signals (Figures 3G-J). These analyses revealed that transcription inhibition and Exosc3 depletion resulted in decreased overlaps of H2B and MED19 signals (Figures 3I-J).

**Figure 3.**
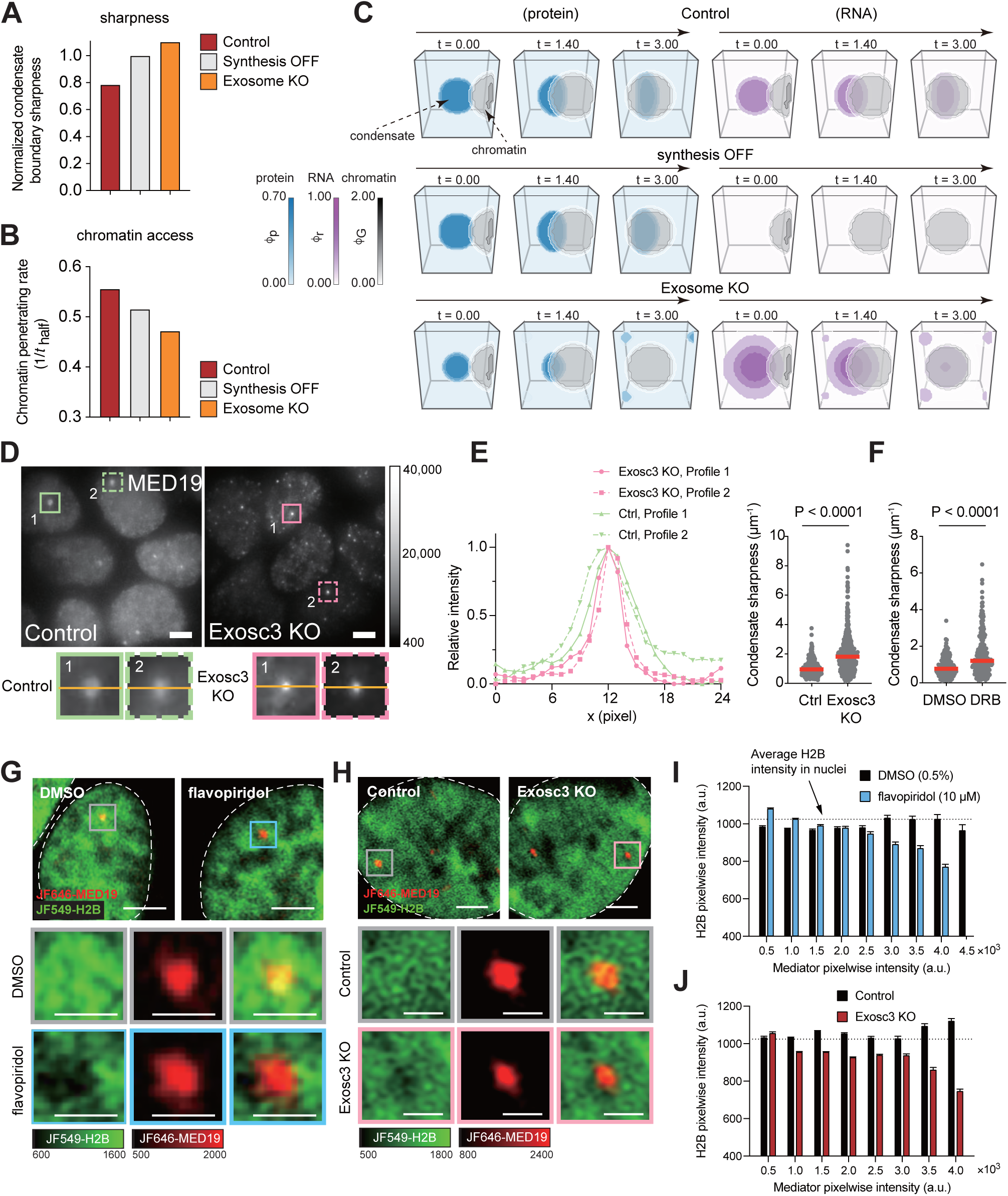
RNA regulation of sharpness and chromatin accessibility of transcriptional condensates. (A, B) Simulation results showing the effects of inhibition of RNA transcription (synthesis OFF) and Exosome KO on the condensate sharpness (A) and chromatin accessibility (B). (C) Visualization of protein (blue), RNA (magenta), and chromatin (gray) concentration fields over simulation time for 3D simulations. The condensate is centered in the field, and the chromatin moves from right to left in the field. (D, E) Changes in TC sharpness. Representative widefield images of Mediator condensates in control and Exosc3-depleted cells (D, top). Solid and dashed boxes indicate TCs, and enlarged images are also shown (D, bottom). Relative intensity profiles of Halo-MED19 signal along the orange lines in the cropped images are shown (E, left). Jitter plots show the changes in sharpness upon Exosc3 depletion (E, right). P value, two-tailed *t*-tests with Welch’s correction. (F) The changes in condensate sharpness upon DRB treatment (20 min), as in (E). (G, H) Halo-MED19 and SNAP-H2B were visualized in cells treated with DMSO or flavopiridol (10 µM, 20 min) (G) or in control and Exosc3-depleted cells (H). Boxes indicate TCs, and enlarged images are shown. Scale bars, 3 µm (overall) and 1 µm (enlarged). (I, J) Bar plots illustrating the relationship between pixelwise intensities of H2B and MED19 in cells treated with DMSO or flavopiridol (10 µM, 20 min) (I) or in control and Exosc3-depleted cells (J). See also Figure S3.

### Loss of RNA exosome reduces Mediator binding at SEs and alters transcriptional pausing

The physical properties of biomolecular condensates—such as viscosity, surface tension, and molecular exchange kinetics—are closely linked to their cellular functions.^45–48^ TCs are involved in regulating the transcription of SE-associated genes^23–25,27^; thus, we next sought to define the functional aspects of gene regulation upon loss of the RNA exosome. To obtain a global landscape of epigenetic and transcriptional changes upon RNA exosome depletion, we performed multi-omics analyses: (1) ChIP-seq (chromatin immunoprecipitation followed by sequencing) for binding of Mediator (MED1), Pol II, and pausing regulators DSIF and NELF, (2) PRO-seq (precision run-on sequencing) for transcription activity and pausing, and (3) RNA-seq for gene expression (Figures 4A-B and S4A). After identifying SEs in our cell lines, which largely overlapped with the previous study (Figure S4B),^23^ we found that RNA exosome depletion results in substantial reduction in MED1 occupancies at SEs, which were more sensitive than typical enhancers (TEs) (Figures 4C and S4C). Albeit relatively modest, Pol II occupancies showed similar trends (Figures 4D and S4C). The high sensitivity of SEs relative to TEs was reminiscent of the effects of depletion of master TFs, Mediator and Brd4, and the effects of 1,6-hexanediol.^6,23,29^ PRO-seq analysis next validated that eRNA transcription was more active in SEs than in TEs and preferentially repressed at SEs upon Exosoc3 depletion (Figure 4E). Based on *de novo* transcriptome assembly using RNA-seq datasets (Figure S4D), we compared transcriptional status and expression levels of various non-coding RNAs, assessed by PRO-seq and RNA-seq, respectively. Loss of RNA exosome preferentially reduced transcription of SE-derived eRNAs (seRNAs) and uaRNAs in PRO-seq analysis, while causing substantial RNA accumulation in RNA-seq analysis (Figure 4F).

**Figure 4.**
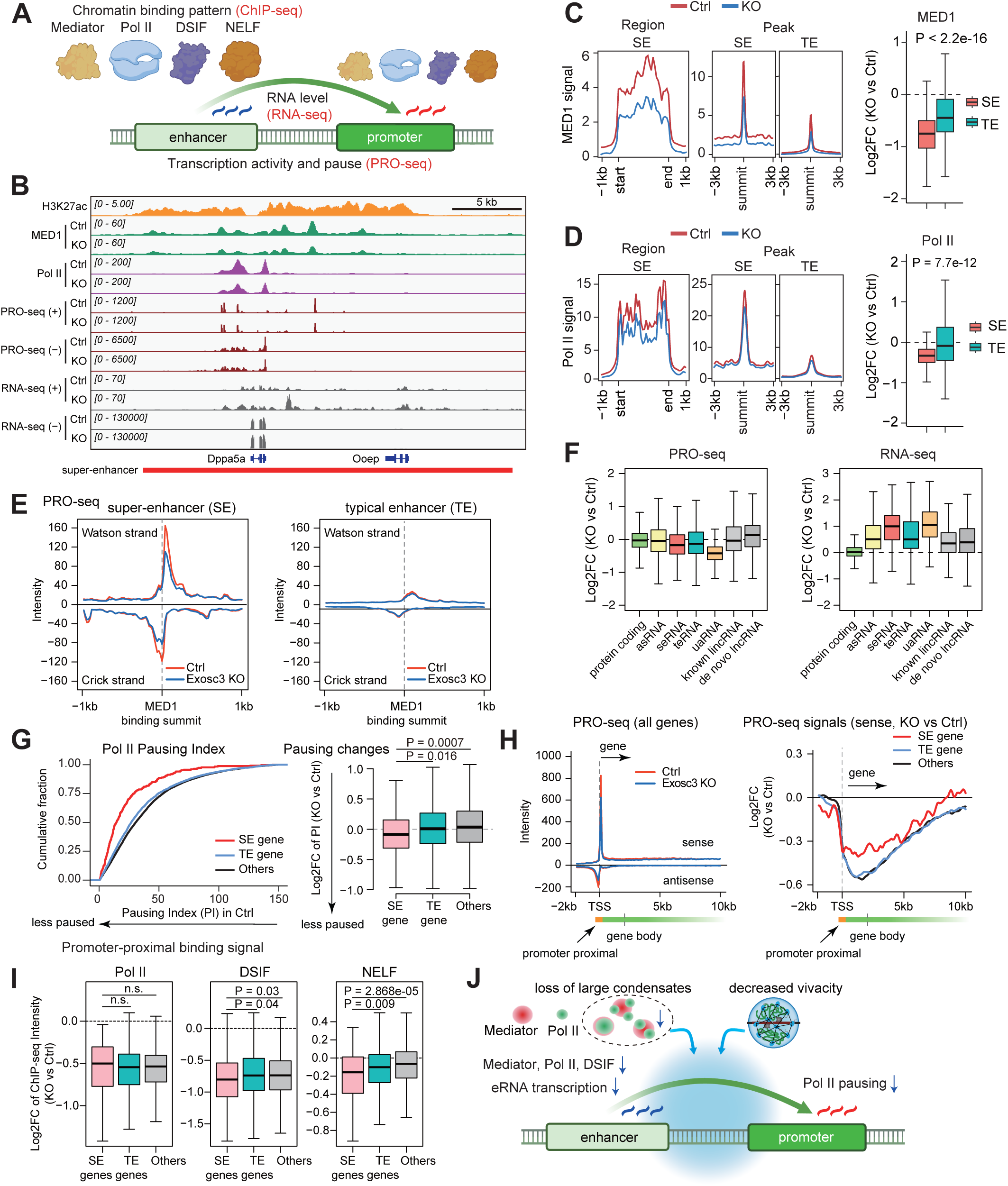
Loss of RNA exosome reduces Mediator binding at SEs and alters transcriptional cycles. (A) Experimental approaches used to investigate chromatin binding patterns of Mediator, Pol II, DSIF, and NELF, RNA expression from enhancers and promoters, and transcriptional activity and pausing. (B) Representative IGV snapshots showing ChIP-seq, PRO-seq, and RNA-seq signals of control and Exosc3-depleted cells around SE and the SE-nearest gene (*Dppa5a*). The H3K27ac ChIP-seq data is from *Yang* et al., 2019.^107^ (C, D) Metaplots displaying mean input-subtracted MED1 (C) and Pol II (D) ChIP-seq signal intensities across SE regions and around SE and TE constituents (left). Boxplots represent the log2-fold change (Exosc3 KO vs Ctrl) in ChIP-seq signals within SE and TE regions (right). P value, Wilcoxon rank sum test. (E) Metaplots of the mean PRO-seq signals in ±1 kb window centered on enhancer constituents for SEs and TEs. (F) Boxplots illustrating log2-fold change (Exosc3 KO vs Ctrl) in PRO-seq (left) and RNA-seq (right) signals for different transcript types. asRNA (antisense RNA), seRNA (super-enhancer RNA), teRNA (typical enhancer RNA), and uaRNA (upstream antisense RNA). (G) Pol II pausing analysis. CDF plots showing the pausing index (STAR Methods) of SE-nearest, TE-nearest, and other genes, as estimated from PRO-seq data (left). Boxplots display the log2-fold change (Exosc3 KO vs Ctrl) of pausing index (right). P value, Wilcoxon rank sum test. (H) Metaplots illustrating PRO-seq signal within −2 kb to +10 kb regions surrounding the TSSs (left) and the log2-fold change (Exosc3 KO vs Ctrl) (right). P value, Wilcoxon rank sum test. (I) Boxplots showing the log2-fold change (Exosc3 KO vs Ctrl) in promoter-proximal ChIP-seq signals for Pol II, DSIF, and NELF in SE-nearest, TE-nearest, and other genes. P value, Wilcoxon rank sum test. (J) A schematic summarizing changes in TC vivacity, SEs, and genes under Exosc3 depletion (Figures 1-4). Images created in BioRender were used in (A) and (J). See also Figure S4.

We further examined how Exosc3 depletion affects transcription of SE-associated genes by examining the status of Pol II pausing and pausing factor loading. As previously reported,^49^ SE genes exhibit lower pausing indices than other genes, indicating efficient pause-release (Figure 4G, left). The pausing indices became even smaller upon Exosc3 depletion, with a significantly greater effect on SE-associated genes compared to other genes (Figure 4G, right). Metaplot analysis surrounding transcription start site (TSS) revealed that a similar reduction in peaks around the pause site for SE-associated genes and other genes, whereas the reduction in gene body signals was milder in SE-associated genes (Figure 4H), explaining the differences in pausing index changes. ChIP-seq analyses of DSIF and NELF, both of which are critical for Pol II pause-release,^3,5^ uncovered that their occupancy in promoter-proximal regions decreased for all gene types after Exosc3 depletion, with the strongest reduction observed in SE genes (Figure 4I). As for Pol II, no significant differences between SE-associated genes and other genes were observed (Figure 4I).

In addition, consistent with the previous report,^50^ we observed that DSIF and NELF are enriched at SEs, relative to TEs (Figures S4E-F). In Exosc3-depleted cells, DSIF occupancy decreased significantly at SE regions, while NELF levels remained unchanged (Figures S4E-F). These collectively suggest that loss of large TCs and reduction in intra-condensate protein mobility and chromatin accessibility (thus reduced locus-targeting probability) coincide with reduced accumulation of Mediator, Pol II, and DSIF at SEs, reduced eRNA transcription, and decreased Pol II pausing at the cognate promoters with reduced DSIF and NELF signals (Figure 4J).

### Increased transcriptional noise of SE-associated genes upon TC perturbations

Despite substantial epigenetic and transcriptional changes around SEs, we failed to detect preferential downregulation of expression levels of SE-associated genes in PRO-seq and RNA-seq analyses (Figure S4G), which were observed upon silencing of Mediator and master TFs.^6^ However, this is ostensibly consistent with no morphological changes in Exosc3-depleted cells in 2i conditions (Figure S1A), whereas they undergo differentiation phenotypes in FBS/LIF conditions.^43^ We reasoned that selection of undifferentiated cells by 2i conditions may contribute to little changes in bulk PRO-seq and RNA-seq and mask fluctuations in gene expression at single-cell level.^51^ Given that Pol II-paused genes have lower cell-to-cell expression variability,^52–55^ we next tested whether disrupted TC homeostasis in Exosc3-depleted cells perturbs cell-to-cell variability of gene expression (Figure 5A). By performing single-cell transcriptome analysis (3′-scRNA-seq analysis), we first confirmed that the vast majority of Exosc3-depleted cells kept a naive ground state in 2i conditions (Figures 5B and S5A). On the other hand, in FBS/LIF conditions, mESCs transitioned into a primed/differentiated-like state upon Exosc3 depletion, whose trajectory was distinct from differentiation with LIF withdrawal, reflecting distinct transcriptome changes (Figures 5B and S5A). Consistent with previous reports,^56,57^ higher transcriptional heterogeneity in FBS/LIF conditions relative to 2i conditions was confirmed.

**Figure 5.**
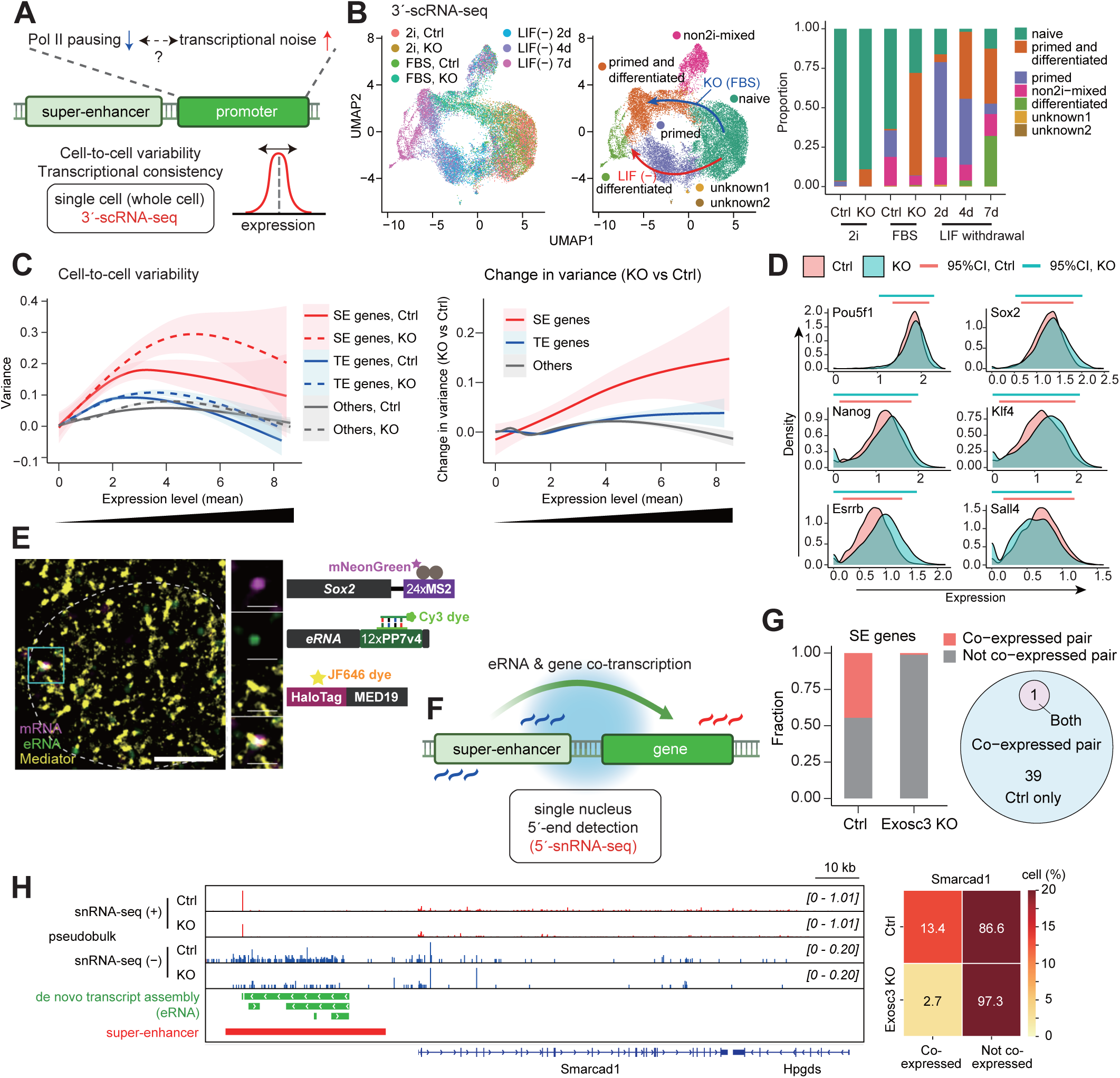
RNA exosome is required for suppressing transcriptional noise and empowering eRNA-gene co-transcription of SE-associated genes. (A) The decrease in Pol II pausing upon Exosc3 depletion can cause the increase in transcriptional noise (See main text for details). Cell-to-cell variability of gene expression was analyzed using 3′-scRNA-seq. (B) UMAP of integrated 3′-scRNA-seq data from mESCs cultured under the indicated conditions. The left panel is colored by culture conditions, while the middle panel represents the pluripotency and differentiation status, with red and blue arrows indicating two differentiation trajectories. The right bar plots show cell state proportions shown in the middle panel. (C) Generalized Additive Model (GAM) smoothed line plots illustrating the relationship between expression level and biological expression variance (left) and the relationship between expression level and changes in biological variance (right). Shaded areas indicate the 95% confidence intervals of the GAM fits. (D) Density plots illustrating the biological variability of expression levels of representative pluripotency genes in control and Exosc3-depleted cells. Top bars indicate the range of 95% of the distribution around the median. (E) A representative image showing Sox2 gene and eRNA co-bursting around MED19 clusters. Scale bar, 3 µm; cropped and zoomed images, 1 µm^2^ area centering the bursting site. (F) Co-transcription of eRNAs transcribed from SE regions and their nearest gene was analyzed using 5′-snRNA-seq. (G) Bar plots showing the fraction of SE genes co-expressed with eRNA from the cognate SEs (left). Co-expressed pairs were further categorized as “Ctrl only”, “Both”, and “KO only” in the Venn diagram (right). No pairs were exclusive to the KO condition. (H) Representative IGV snapshots showing pseudobulk 5′-snRNA-seq signals in control and Exosc3-depleted cells around SE and the SE-nearest gene (*Smarcad1*). The right heatmap illustrates the proportion of gene-expressing cells that either co-expressed eRNAs or did not. Images created in BioRender were used in (A) and (F). See also Figure S5.

We next calculated biological variance for each gene to measure cell-to-cell variability and transcription noise (Figures 5C-D and S5B-C) (STAR Methods). In agreement with the previous study,^58^ and as can be inferred from the lower Pol II pausing (Figures 4G-H), SE-associated genes exhibited higher variance than other types of genes (Figure 5C). Strikingly, Exosc3 depletion led to further increase in variance for SE-associated genes, preferentially affecting genes with moderate to high expression levels (Figures 5C and S5B). These trends were confirmed for key pluripotency genes (Figure 5D). In contrast, non-SE-associated genes showed no significant alterations in cell-to-cell variability (Figures 5C and S5B). Consistent with the results of bulk analyses, changes in expression levels did not differ between SE-associated and TE-associated genes (Figure S5C). These results suggest that the loss of normal TCs—caused by elevated eRNA levels—disrupts SE functionality, ultimately leading to heterogenous transcription of SE-associated genes.

### Impaired eRNA-mRNA co-expression upon RNA exosome depletion

TCs are thought to physically incorporate SE and the corresponding promoter to drive transcription.^23,24,28^ Recent single-cell analyses have highlighted the concurrent transcription of enhancer-gene pair.^59–61^ In fact, we observed that eRNA and mRNA signals at Sox2 loci correlated well and frequently colocalized with Mediator TCs (Figures 5E and S5D). To study the impacts of Exosc3 depletion on eRNA-gene coordination, we next performed 5′-single-nucleus RNA-seq (5′-snRNA-seq), which can reflect transcription status more directly relative to 3′-scRNA-seq (Figures 5F-H and S5E-F). We quantitated eRNA signals using our *de novo* transcriptome assembly results (Figure S4D). As previously described,^62–64^ 5′-snRNA-seq successfully captured eRNA signals that corresponded well to eRNA regions (Figures 5H and S5E). We identified 90 SE-gene pairs, in which both eRNA and gene expression were detected in both control and Exosc3-depleted cells, and then determined co-expression status using pairwise Pearson correlation values and chi-square tests. As a result, 39 co-expressed pairs (43.3%) were identified in control cells, whereas only 1 pair (0.01%) remained co-expressed in Exosc3-depleted cells (Figure 5G). This strongly suggests a loss of co-transcription of eRNA-gene pairs at SEs following Exosc3 depletion. Figures 5H and S5E-F show representative examples of co-expressed eRNA-gene pairs at SEs.

### RNA exosome-dependent widespread contacts between distant SE-containing A compartments

SEs are thought to activate the corresponding neighborhood genes in the local chromosomal structures, called insulated neighborhoods, within a single TAD, which keeps the integrity of local gene regulation.^15^ Besides, inter-TAD associations, which involve multiple SE regions, are frequently observed. These are called TAD triplets in previous reports.^19,20^ These genome organizations can be partly recognized as a manifestation of TC and chromatin interactions. Therefore, we next analyzed Hi-C data to investigate how loss of large TCs and alterations in TC qualities upon Exosc3 depletion correlate with 3D genome organization (Figure 6). As shown in Figure 6A, positions of A compartments (regions with positive PC1 values) corresponded well to the regions with high MED1, high H3K27ac, and low H3K9me2 signals. The distribution of euchromatin compartment (A compartment) and heterochromatin compartment (B compartment) did not largely change upon Exosc3 loss (Figure 6B). Importantly, by inspecting the gross appearance of contact changes across the entire chromosome, we found that RNA exosome depletion led to a widespread decrease in contacts between A compartments, which contain SEs (Figure 6B). These changes were observed in most of the chromosomes (Figures S6A-D). Nearly all SE-containing compartments belonged to A compartments, whereas TE-containing and enhancer-free compartments were evenly distributed between A and B compartments (Figure 6C). SE-containing A compartments (SE-A compartments) differ from TE-containing A compartments (TE-A compartments) in several aspects: SE-A compartments are longer (Figure S6E) and more enriched with MED1 (Figure S6F), whose occupancies are more drastically reduced following Exosc3 depletion (Figure 6D).

**Figure 6.**
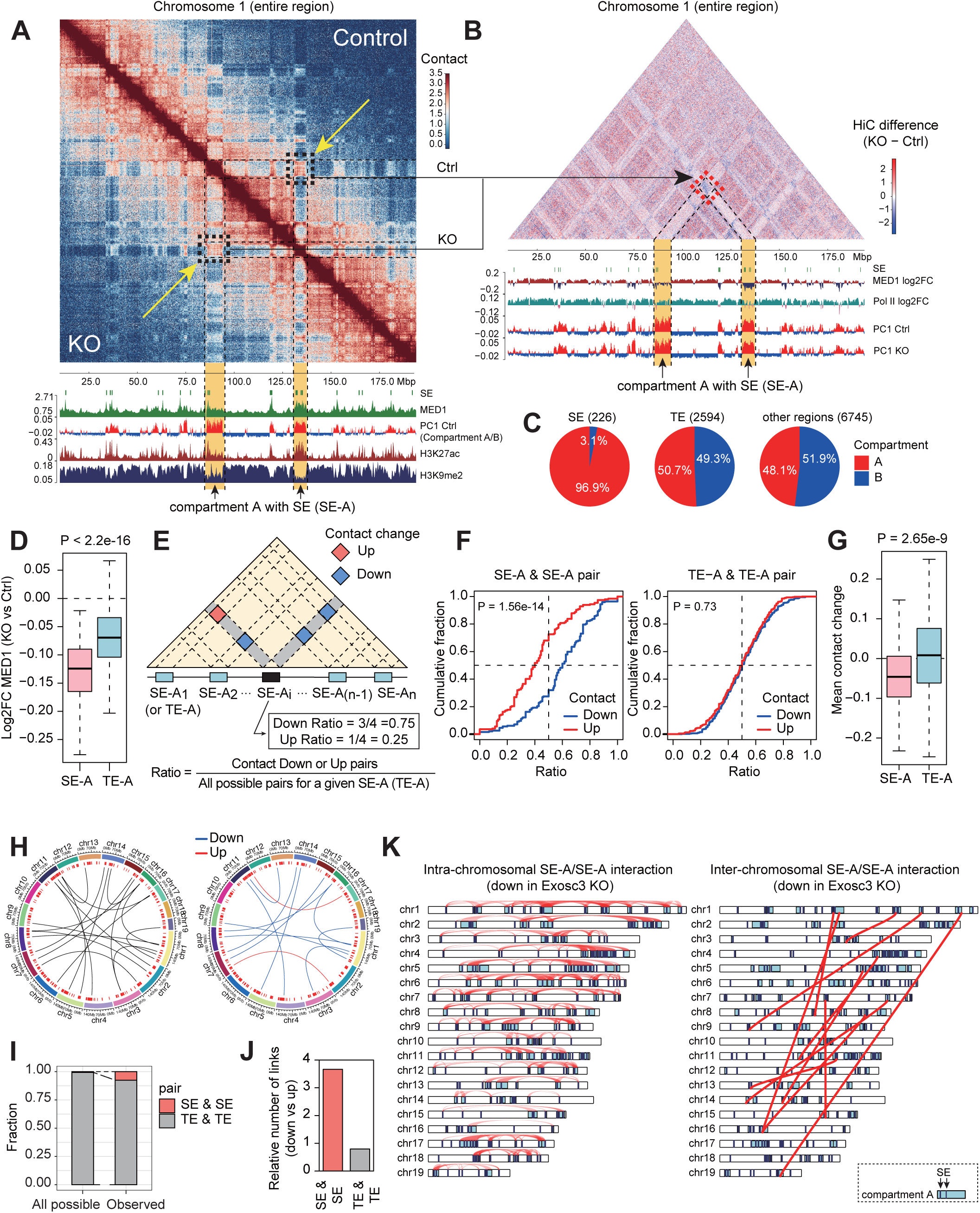
Widespread interactions between SE-containing A compartments and the perturbations upon RNA exosome loss. (A) Representative Hi-C results showing contact matrices of entire chromosome 1 at 200 kb resolution in control and Exosc3 KO conditions. Coverage tracks of ChIP-seq signals of MED1 (control), H3K27ac, H3K9me3, and PC1 scores (compartment A/B), as well as the position of SEs are shown below. Matrices are KR normalized. H3K27ac and H3K9me3 ChIP-seq data are from *Yang et al.*, 2019.^107^ (B) Differential contact matrices showing interaction changes after Exosc3 KO. Bottom tracks display the log2 FC (KO vs Ctrl) of MED1 and Pol II ChIP-seq signals and PC1 scores in Ctrl and Exosc3-depleted cells. (C) Pie charts illustrating the percentage of A and B compartments associated either with SEs or TEs. The proportions of enhancer-free compartments A and B (other regions) are also shown. (D) Boxplots showing the log2-fold change (Exosc3 KO vs Ctrl) in mean MED1 ChIP-seq signal density. P value, Wilcoxon rank sum test. (E) A diagram depicting contact changes between SE-or TE-containing A compartments (SE-A or TE-A) and how the ratio was calculated as the proportion of “Down” or “Up” pairs to all possible pairs for a given compartment. (F) CDF plots of the “Down” or “Up” ratio for SE-A/SE-A pairs involving each SE-A (left) and TE-A/TE-A pairs involving each TE-A (right). P value, Kolmogorov-Smirnov test. (G) Boxplots showing the distribution of mean contact changes for each compartment of SE-A and TE-A (> 0.2 Mb) (KO vs Ctrl). P value, Wilcoxon rank sum test. (H) Circular diagrams of inter-chromosomal interactions. All the significantly changed interactions following Exosc3 depletion (left), and the “Up” (contact changes > 0.2) and “Down” (contact changes < −0.2) interactions (right) are shown. Red bars indicate SE-A compartments. (I) Fractions of SE-SE and TE-TE compartment pairs calculated for all possible combinations and observed interactions. (J) Bar plots showing the relative numbers of altered links (Down vs Up) in SE-SE and TE-TE pairs. (K) Karyoplots with links showing intra- and inter-chromosomal interactions between two SE-containing A compartments where contacts are decreased after Exosc3 KO. See also Figure S6.

To generalize our findings regarding contacts between SE-A compartments, we next performed pairwise computation of the mean contact changes for SE-A compartment pairs and TE-A compartment pairs on the same chromosome (STAR methods). We then calculated the ratio of the number of contact-decrease pairs (or contact-increase pairs) relative to the total number of possible pairs per compartment (Figures 6E and S6G). Remarkably, SE-A compartments were skewed toward decreased contacts in the absence of Exosc3, while TE-A compartments showed no significant differences (Figures 6F and S6G). Accordingly, mean contact changes for each SE-A and TE-A compartment indicate that inter-SE-A compartment interactions, but not inter-TE-A compartment interactions, are prone to decrease upon Exosc3 loss (Figure 6G).

We further analyzed inter-chromosomal interactions (Figures 6H-J). In total, we identified 21 SE-SE and 257 TE-TE interacting compartment pairs across chromosomes, and found that the number of SE-SE interactions were significantly more frequent than TE-TE interactions, when compared to expected frequency (Figure 6I, Fisher’s exact test P = 4.0e-21, Odds ratio = 30.3). Notably, SE-SE interactions tended to decrease, whereas TE-TE interactions showed no directional bias (Figure 6J). Taken together, these findings indicate the presence of RNA exosome-dependent widespread inter- and intra-chromosomal interactions of SE-A compartments (Figures 6K and S6H), echoing with the condensate vivacity-dependent chromatin accessibility (Figure 3).

### Globally coordinated transcription of SE-associated genes

Although SE-involved high-order interactions like TAD triples are frequently observed in recent genome organization studies,^19,20,65,66^ their functional significance is currently obscure. We speculated that multiple SE-associated genes, which interact with TCs and associate with SE-A compartment interactions, may exhibit coordinated transcriptional bursts (Figure 7A). We therefore performed live-cell imaging to examine co-transcription of two gene pairs; Sox2-Tet2 and Nanog-Smarcad1—both SE-associated pluripotency genes located in a pair of distant but interacting A compartments (Figures 7B-C and S7A-B, and Videos S3 and S4). Consistent with our hypothesis, we detected spatio-temporal proximity of transcriptional bursts of two genes, while two genes are 98.9 Mb apart on chromosome 3 in the case of Sox2-Tet2 pair (Figures 7B-C and S7A). The timing of transcriptional bursts from Sox2 and Tet2 genes largely coincides, and both transcription sites transiently associate with a TC. We further examined this possibility by utilizing our 5′-snRNA-seq datasets. To do so, we systematically analyzed gene-gene co-expression status for 97,853,055 pairs (13,990 genes) by calculating Spearman correlation coefficients, associated P values, and additional permutation-based P values (Figure 7D) (STAR methods). We extracted significantly co-expressed gene pairs, using the following criteria: Spearman’s r > 0.1, adjusted P value < 0.01, and adjusted P value from permutation test < 0.01 (Figure 7E). Spearman correlation coefficients of SE-associated gene pairs were higher than those for pairs of TE-associated genes and all gene pairs (Figure S7C). Furthermore, the fraction of significant co-expression pairs of SE-associated genes was markedly higher, exceeding those of TE-associated gene pairs and randomly selected gene pairs (Figure 7E). Correlation heatmaps of SE-associated genes revealed large co-expression networks in control cells (Figure 7F). Strikingly, the overall correlation coefficients were substantially decreased in Exosc3-depleted cells, although the overall cluster pattern was partially retained (Figures 7F and S7C). Intriguingly, intersection with eRNA-gene co-expression status and Hi-C contact changes further revealed that the co-expression network (Figure 7F, cluster A) was enriched with eRNA-gene co-expressing pairs (Figure 7G) and Exosc3-dependent SE-A interactions (Figure 7H). Notably, correlation heatmaps derived from 3′-scRNA-seq data failed to detect this co-expression pattern and changes associated with RNA exosome loss (Figure S7D), suggesting that global co-transcription is masked by cytoplasmic mRNAs.

**Figure 7.**
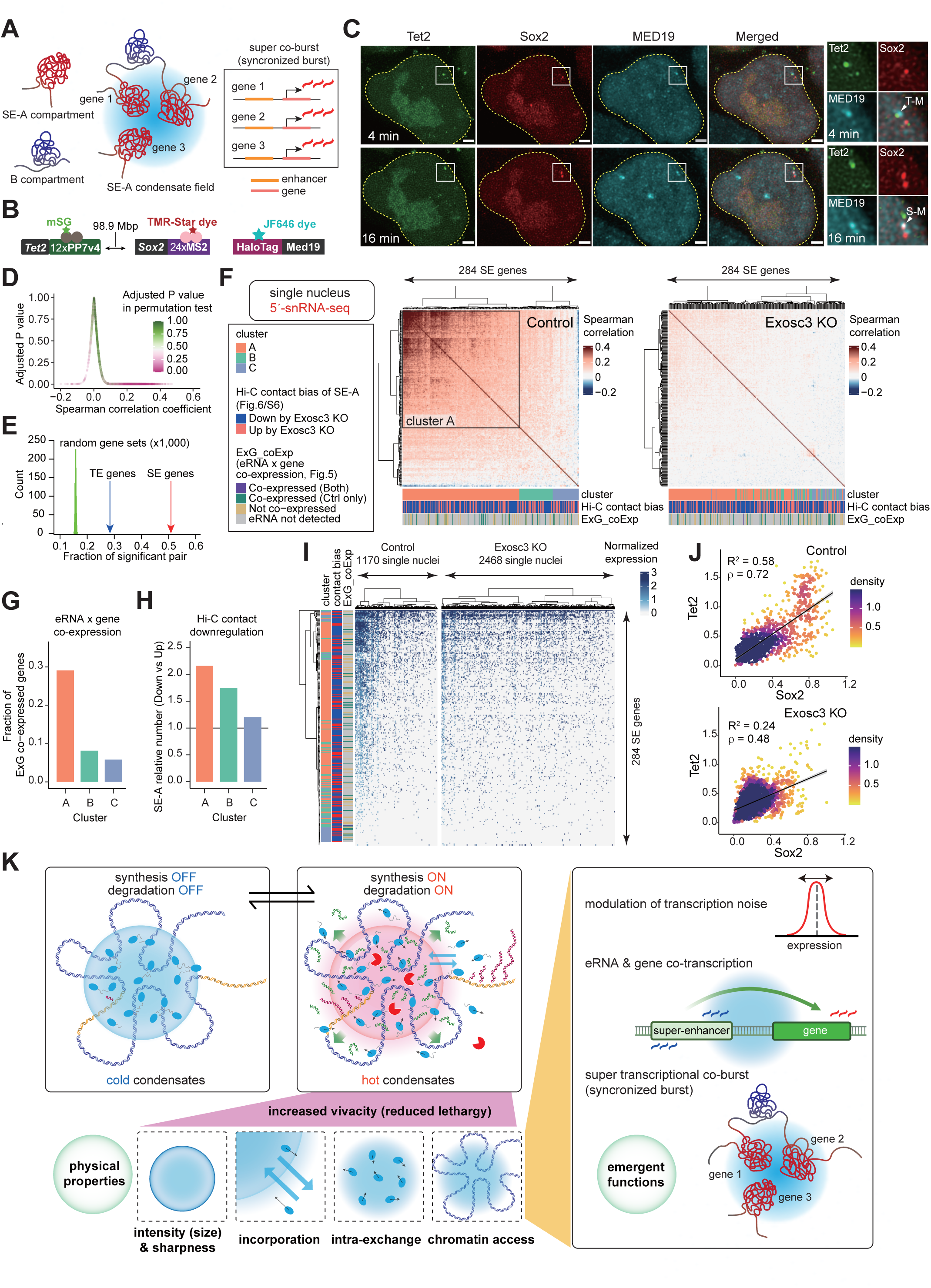
SE-associated genes are transcribed in a synchronized manner that are sensitive to RNA and condensate perturbations. (A) A schematic of widespread synchronized transcription of SE genes in the SE-containing A compartments (SE-A). Vivacious transcriptional condensates are presumed to incorporate multiple SE-As and drive synchronized transcriptional bursting. Consequently, higher gene-gene expression correlations are expected among SE-associated genes. (B) A schematic showing our live-cell imaging system. Sox2 and Tet2 genes are 98.9 Mbp apart. (C) Representative live-cell images showing co-bursting of endogenous *Sox2* and *Tet2* genes in a control cell. We observed substantial temporal overlap between Tet2 and Sox2 bursts (Tet2: 0-4 and 16-24 min, Sox2: 4-24 min). Boxes indicate bursts at 4 and 16 min of measurements. The enlarged images show co-localization of Tet2 burst and Mediator condensate at 4 min (arrowhead, T-M) and co-localization of Sox2 burst and Mediator condensate at 16 min (arrowhead, S-M), respectively. Scale bar, 2 µm. (D) Gene-gene co-transcription analysis in 5′-snRNA-seq. Pairwise Spearman correlation coefficients (x-axis), their adjusted P values (y-axis), and adjusted P values in an additional permutation test (color scale) were calculated for all gene-gene pairs in control cells (STAR methods). (E) Quantification of fraction of significantly co-transcribed gene-gene pairs for SE gene pairs (red), TE gene pairs (blue), and 1,000 permutation sets of random gene pairs (green histogram). (F) Heatmaps showing Spearman’s correlation coefficients among SE genes in control (left) and Exosc3-depleted cells (right), calculated using 5′-snRNA-seq data. Clusters A-C were defined based on the Ctrl data (left), and transferred to Exosc3 KO data (right). Hi-C contact bias indicates the tendency of contact changes between compartments where the genes are located (Figures 6E-F). The eRNA-gene co-expression status (ExG_coExp) from Figures 5F-G is also shown. (G, H) Bar plots showing the fraction of eRNA-gene co-expressed genes relative to the number of genes where the expression of their associated eRNAs detected in 5′-snRNA-seq (G) and the number of “Down” compartments relative to “Up” compartments (H) in clusters A-C defined in (F). (I) Heatmaps displaying sqrt-normalized counts of each gene per cell, calculated from 5′-snRNA-seq data. Clusters A-C, Hi-C contact bias, and eRNA-gene co-expression information are as in (F). (J) Correlations of single cell expression levels of two SE genes, Nanog and Smarcad1, in 5′-snRNA-seq. Lines represent linear regression with 95% confidence intervals (shaded areas). *R*^2^, coefficient of determination; ρ represents Spearman’s correlation coefficient. (K) A summary model describing RNA-mediated regulation of TC vivacity and the functional impacts of “hot” condensates on coordinated transcription. See the Discussion for details. Images created in BioRender was used in (K). See also Figure S7.

We also found that SE-associated co-expression networks were prominent in a subset of control cells (approximately ∼25%) (Figure 7I), reminiscent of the observation that ∼40% of the E3.5 inner cell mass cell nuclei showed RNA-FISH signals for pluripotency genes such as *Nanog*.^67^ This co-expressing subset was largely diminished upon Exosc3 loss (Figure 7I). Given that burst durations of Sox2-Tet2 and Nanog-Smarcad1 pairs substantially overlap in live-cell imaging (Videos S3 and S4), these co-expression networks in 5′-snRNA-seq presumably represent a set of temporally correlated co-transcription. Representation of individual pair of SE-associated genes conspicuously demonstrated the presence of a subset of cells that exhibit strong bursting activities of two genes simultaneously in an RNA exosome-dependent fashion (Figures 7J and S7E-J). In summary, these findings highlight that SE-associated genes, residing in Exosc3-dependent A compartments and presumably interacting with normal TCs, form coordinated co-burst transcription networks, which are disrupted upon Exosc3 depletion.

## DISCUSSION

Our findings presented here suggest an elaborated RNA-mediated condensate model in gene control, where both synthesis and decay of short RNAs from enhancers optimize TC formation, and TCs do not merely activate transcription but coordinate transcriptional consistency of multiple genes. We envision that TC interconverts between “hot” (vivacious) and “cold” (lethargic) states in response to RNA regulation, with hot condensates being physically dynamic and transcriptionally active, whereas cold condensates are physically static and transcriptionally inactive (Figure 7K). This model harmonizes the positive roles of RNA synthesis and RNA exosome in transcriptional activation, as reported in previous studies: RNA molecules facilitate Mediator recruitment and TF trapping and stimulate Pol II pause release,^68–71^ while RNA exosome prevents accumulation of deleterious R-loop structures for proper enhancer activities.^39^ Therefore, the elaborated RNA feedback model provides an explanation of why eRNAs are pervasively transcribed but rapidly degraded. Our study indicates that RNA-dependent vivacious TCs represent the key molecular ecosystem to maintain cell identity and phenotypes.

We propose that hot condensates achieve transcription coordination through their fluid and dynamic properties. TC formation is driven by a complex coacervation^72^ process promoted by oppositely charged polymers –proteins acting as polycations and RNAs as polyanions.^37^ Previous studies have demonstrated that RNA molecules are key determinants of condensate physical-biochemical properties, such as viscoelasticity and surface tension,^73,74^ which influence the rate of coalescence. In addition, we observed increased TC boundary sharpness and delayed recovery from photobleaching upon Exosc3 depletion. These changes may reflect a “hardened” interface and decreased molecular exchange kinetics of condensates.^75^ Theoretically, increased interface resistance can arise from (i) the unavailability of sticky residues due to their conformational burial or the saturation of sticker bonds or (ii) the formation of charged layers at the interface, leading to electrostatic repulsion. Excess eRNAs may saturate sticker bonds or form negatively charged layers at the surface, thereby disrupting the formation of larger TCs^76^: meanwhile, the RNA efflux reshapes the free energy landscape of condensates,^77^ creating a steepened gradient that accelerates axial molecular exchanges.

In our experiments, mESCs maintained pluripotent states and exhibited reversible TC changes upon Exosc3 depletion, providing a window to experimentally link changes in TC homeostasis, SE activities, and coordinated transcription. These phenotypes are contrasting to the differentiation phenotypes and DNA damage responses observed in FBS/LIF conditions in our previous report.^43^ These differences can be explained by the effects of 2i condition on cell status, to select undifferentiated cells and maintain high Nanog levels,^51^ and cell cycle effects, to active p53 and elongate G1 phase,^78^ which potentially allows recovery from DNA damage in response to Exosc3 loss. In addition, reversible effects on Exosc3 depletion on TC homeostasis is contrasting to those of 1,6-hexanediol (1,6-Hex). Although 1,6-Hex has been widely employed to dissolve biomolecular condensates, 1,6-Hex impairs kinase and phosphatase activities,^79^ immobilizes and condenses chromatin,^80^ and irreversibly disrupts compartment patterning due to A-B compartment mixing,^81,82^ leading to cell shrinkage and the formation of aberrant aggregates.^81^ These effects can distort the functional interpretation of the condensate changes. In this context, our strategy provides a milder and more suitable approach for TC research, which has depended on the use of 1,6-Hex or the small-scale datasets obtained at one or a few genomic loci, although the secondary effects of Exosc3 depletion are not completely excluded.

Previous studies of SE deletions have established activation roles of SEs in transcription of cell identity genes. On the other hand, while maintaining the overall expression of SE-associated genes, Exosc3 depletion disrupted multiple aspects of coordinated transcription: suppression of transcription noise, eRNA-mRNA co-expression, and gene-gene co-expression associated with pervasive intrachromosomal interactions. These findings suggest that basal transcriptional activation by SE-associated genes and coordinated transcription by vivacious TCs are separable to some extent. Such independency can be partly explained by the differential requirement of structured domains and IDRs of transcriptional regulators for transcriptional initiation and condensate formation, respectively. Supporting this, transcription activation by the Mediator complex is a stoichiometric process, primarily involving structured domains that support the interaction with pre-initiation complex to initiate transcription.^83^ In contrast, the TC formation is driven by non-stoichiometric multivalent interactions between IDRs.^84^ We suspect that the Mediator complex concentration in TCs exceeds the saturation threshold required for Mediator complex-dependent transcription initiation. Even in the absence of normal-sized TCs, a sufficient amount of Mediator may still be recruited to SE constituents to promote efficient transcription initiation. These considerations are largely consistent with the previous reports showing distinct behavior of Pol II clustering and pause regulation at SE-associated and other genes and pliable roles of non-essential Mediator subunits increasing structural complexity of the tail module.^85–87^

Our integrated analyses of 5′-snRNA-seq and Hi-C datasets (Figures 6 and 7) indicate that coordinated transcription of multiple SE-associated genes are associated with widespread long-range interactions of distant SE-As, which were consistent with many-body interactions of SEs, such as TAD triplets, detected by several recent technologies such as GAM, SPRITE, and MERFISH.^19,20,65,66^ Together with co-regulation of distant genes by shared enhancers in *Drosophila* (“topological operon”),^30,88^ our results suggest that coordinated transcription of functionally related genes is a prevalent feature in multicellular organisms. On the other hand, depletion of cohesin causes mixing of small Mediator hubs into large clusters and a global increase in aberrant co-bursts.^89,90^ Therefore, coordinated transcription is elaborated by the balance between insulation mechanisms through CTCF and cohesin and the pervasiveness of TCs^91,92^ and could be influenced by heterogeneity and intrinsic variation in spatial genome organization in individual cells.^93,94^ Meanwhile, an open question remains whether specific SE gene combinations are preferentially co-regulated via large TCs or how frequently each combination forms.

Finally, our results provide key insights into how the TCs are advantageous for the entire biological system. Although the basal transcription levels of SE-associated genes can be maintained under disrupted TC homeostasis, the inherent bursting and stochastic nature of transcription could increase the cells not expressing repertories of desired gene set. In this context, regulation of co-bursts by TCs provides a strong foundation to keep the integrity of cell identity and phenotypes. Subsequent mRNA export and stabilization prolong the duration of expression of cell identity genes to efficiently maintain cell identity even in the transcription bursting scheme. Importantly, recent studies show attenuated coordinate gene expression, reduced Pol II pausing, and increased Pol II elongation speed along with aging.^95–97^ These findings are consistent with our concomitant observations of reduced Pol II pausing and increased transcription noise upon Exosc3 depletion. Furthermore, consistent with reports describing that depletion of RNA exosome triggers senescent phenotypes,^98,99^ reduced expression of RNA exosome subunits observed in senescent cells^98^ may collapse suppression of transcription noise via Pol II pausing. Given that many condensates show degenerated properties in pathological conditions, such as aging and neurodegenerative diseases,^100–102^ disruption in TC homeostasis and coordinated transcription may play versatile roles in various diseases including aging and cancer, where the heterogeneity of cell phenotypes is especially important.

### Limitations of the Study

We validated the relationships between RNA exosome, TC homeostasis, and transcription by integrating simulations, imaging, and multi-omics analyses. However, given the various roles of RNA exosome in RNA homeostasis,^99,103^ further studies are essential to directly examine their causal relationships, especially, how microscopically observed TCs and genome interactions in Hi-C analyses correspond, and how the compositions of TCs in Exosc3-depleted cells alter. To do so, it is important to directly analyze the components and interaction maps of endogenous, but not reconstituted, TCs, although recent studies expand the experimental approaches for TCs.^104,105^ The perspective of such structure-function relationships could be further complemented by high-resolution analysis of transcribing Pol II, such as scGRO-seq,^59^ and genome structures, such as Micro-C,^106^ SPRITE,^20^ and GAM.^19^ Our simulation model is based on a diffusion framework, which cannot address the contribution from potential advective flow owing to the RNA efflux. A more tailored model will be essential to simulate the convection and advection that might occur in a “hot” condensate when RNA synthesis, degradation, and motion are at play.

## ACKNOWLEDGEMENTS

We are grateful to Suzuki laboratory members for discussion on the project. We also appreciate Mayumi Fujisawa, Mayumi Yoshimura, Masako Tsuji, and Kaori Tatsumi for technical supports. We thank Division for Medical Research Engineering, Nagoya University Graduate School of Medicine, for the technical support of the usage of Nikon AXR, specifically Eri Yorifuji and Mayumi Furukawa; the Robert A. Swanson (1969) Biotechnology Center at the Koch Institute for Integrative Cancer Research at MIT for technical support, specifically S. Levine and the staff of the BioMicro Center/KI Genomic Core Facility and G. Paradis, M. Jennings, and M. Saturno-Condon of the Flow Cytometry Core Facility; the W.M. Keck Microscopy Facility of Whitehead Institute for the technical support of super-resolution confocal microscopy. We also thank L.D. Lavis (HHMI, Janelia) for the generous gift of the JF dyes, Masaki Kawamata for the CRISPR protocol, and Dig B. Mahat for the PRO-seq protocol. We dedicate this work in memory of Jingzhi Zhu, a research computing specialist at the Koch Institute Integrated Genomics and Bioinformatics Core, who worked tirelessly to ensure that the community had a robust computing infrastructure.

This work was supported by JSPS Grant-in-Aid for Scientific Research (A) (JP24H00614 to H.I.S.) and (C) (JP20K06925 to K.O. and JP22K07210 to S.Y.), and JSPS Home-Returning Researcher Development Research (19K24694 to H.I.S.); AMED (JP22ama221111, JP23ck0106791, and JP23kk0305026 to H. I. S. and JP23tk0124003 to H.I.S. and K.G.N.S.); the Takeda Science Foundation (to K.O., S.Y., and H.I.S.); Toray Science Foundation (22-6304 to H.I.S.); Inamori Research Institute for Science (InaRIS) (to H.I.S.); Program Project Grant P01CA042063 from the National Cancer Institute, United States Public Health Service Grants R01-GM34277 and R01-CA133404 from the National Institutes of Health, and, in part, the Koch Institute Support (Core) Grant P30-CA14051 from the National Cancer Institute (to P.A.S.); JSPS fellowship for Young Scientists (to Y.S.); Nagoya University WISE Program CIBoG and JST-SPRING Program (to Y.S., M.S., and T.M.).

We are profoundly grateful to Richard A. Young (Massachusetts Institute of Technology; Whitehead Institute for Biomedical Research) for his invaluable intellectual input and for his generous support of M.M.Z., provided in part by grants to Richard A. Young: the National Institutes of Health grant (GM144283) and the National Science Foundation grant (PHY2044895).

## AUTHOR CONTRIBUTION

K.Ogami., M.M.Z. and H.I.S. conceptualized the project. K.Ogami., M.M.Z., M.S., T.M., and K.G.N.S. performed imaging analyses. M.M.Z performed simulation analyses. K.Ogami., S.Y., Y.S., K.Onimaru., and S.K. contributed to multi-omics library preparation. K.Ogami. and H.I.S. performed multi-omics computational analyses. K.Ogami., M.M.Z., H.I.S. wrote the original draft, and P.A.S. performed critical review. H.I.S. supervised the project and writing of the manuscript.

## STAR METHODS

### Key Resource Table

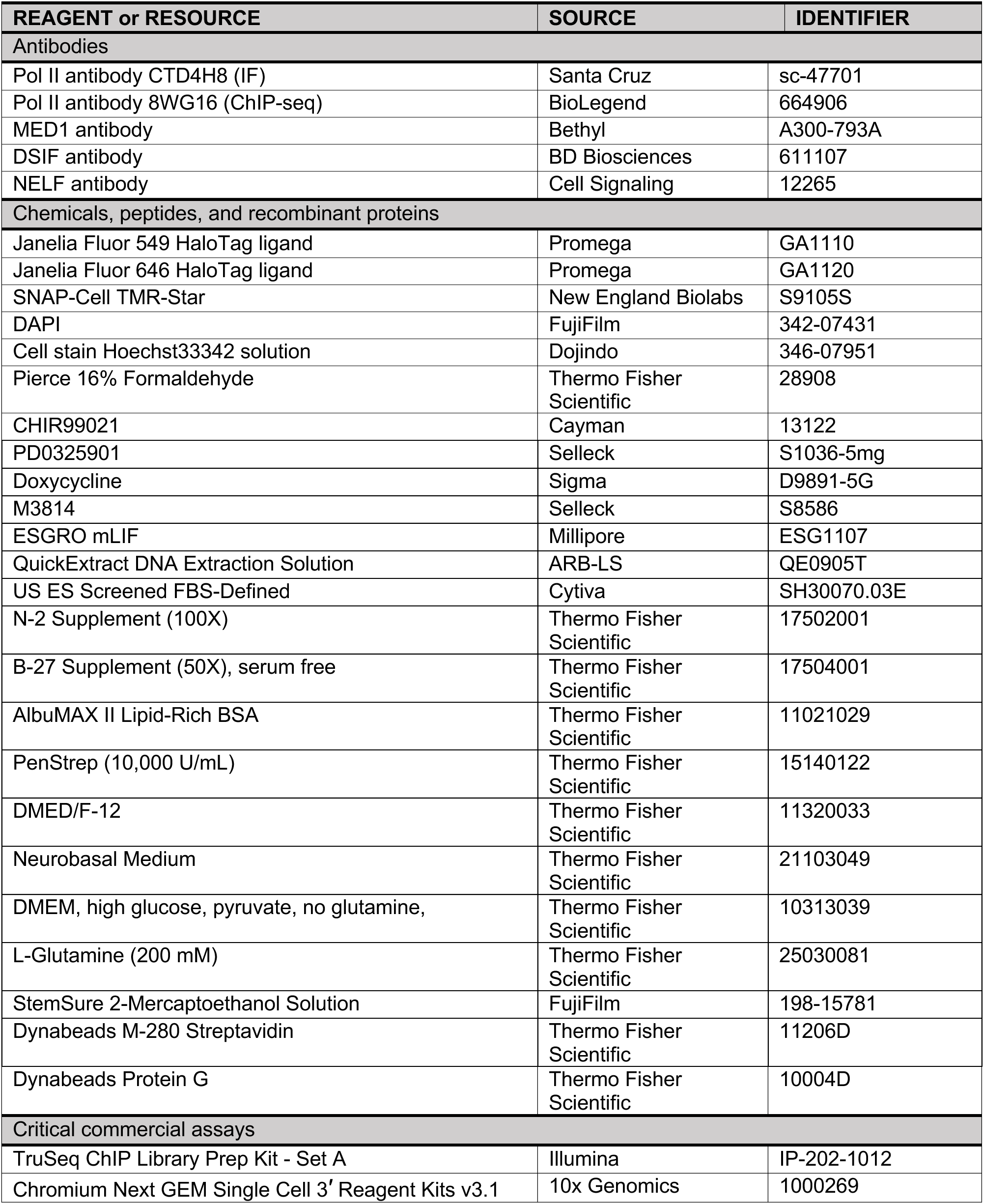
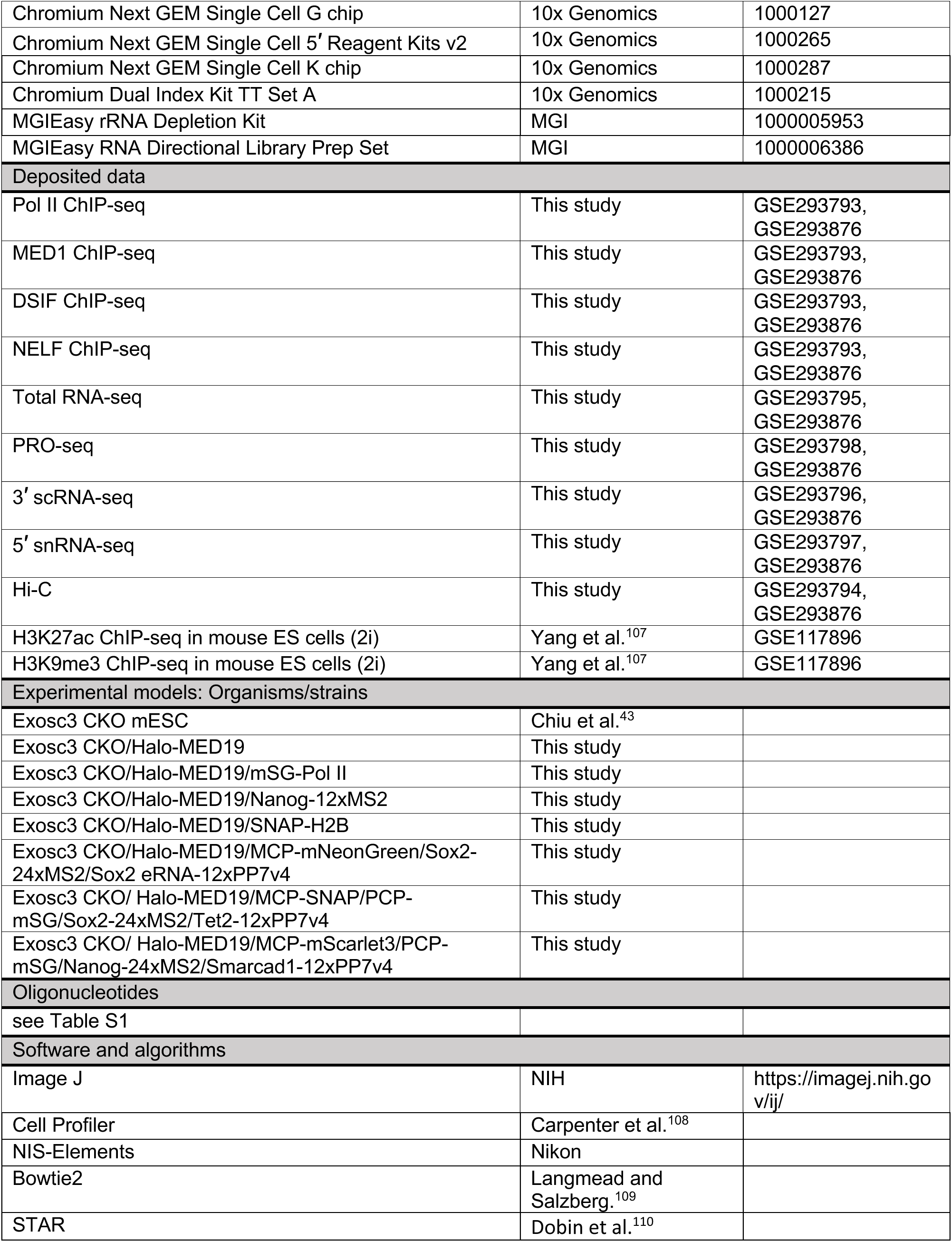
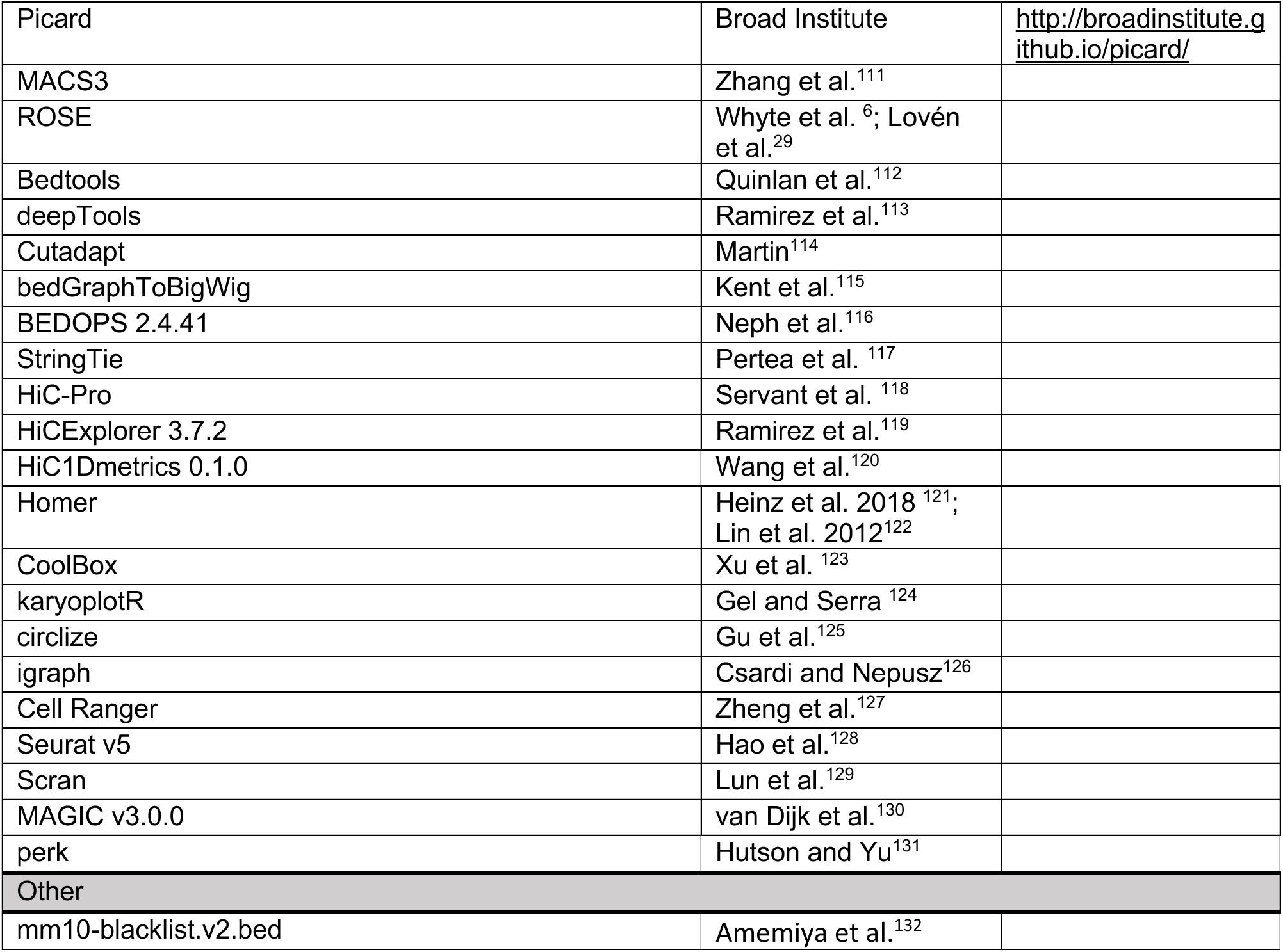

## RESOURCE AVAILABILITY

### Lead Contact

Further information and requests for resources and reagents should be directed to and will be fulfilled by the Lead Contact, Hiroshi I. Suzuki.

### Materials Availability

Materials associated with the paper are available upon request.

### Data and Code Availability

- The accession number for the data for ChIP-seq, PRO-seq, RNA-seq, scRNA-seq, snRNA-seq, and Hi-C reported in this paper is GEO: GSE293876.
- Codes used in this study are available from lead contact upon request.
- Any additional information required to reanalyze the data reported in this paper is available from the lead contact upon request.

## EXPERIMENTAL MODEL AND SUBJECT DETAILS

### Cell Culture

All mESC lines used in this study, except otherwise described, were cultured under serum-free 2i condition without feeder cells (1:1 of DMEM/F-12 and Neurobasal media, supplemented with 1 x N2 supplement, 1 x B27 supplement, 100 U/ml PenStrep, 3 µg/ml AlbuMAX II, 1 x MEM NEAA, 1 mM Sodium pyruvate, 1 mM L-glutamine (all from Thermo Fisher Scientific), 0.1 µM 2-Mercaptoethanol (FujiFilm), 1 mM PD0325910 (Selleck), 3 mM CHIR99021 (Cayman), 0.1 µg/mL Doxycycline (Dox, Sigma), and 100 U/ml mLIF (EMD Millipore). Before seeding, dishes were coated with either 0.2% gelatin for more than 10 minutes at room temperature or 5 µg/ml poly-L-ornithine (PLO, Sigma) in 1 x PBS buffer at 37°C for more than 2 hours, followed by 5 µg/ml Laminin (FujiFilm) in 1 x PBS buffer for more than 2 hours at 37°C. Depending on the experiments, cells were cultured in DMEM containing 10 mM HEPES, 1 x MEM NEAA, 1 mM L-glutamine, 100 U/ml PenStrep (all from Thermo Fisher Scientific), 0.1 µM 2-Mercaptoethanol (FujiFilm), 0.1 µg/mL doxycycline (Sigma), 100 U/ml mLIF (EMD Millipore), and 15% ES certificate FBS (Cytiva, SH30070.03E). Conditional knockout of Exosc3 was achieved by culturing cells in doxycycline-free media for 65 hours after three times wash with PBS (-). Differentiation was promoted by culturing cells under FBS condition without mLIF for 2, 4, and 7 days. In all the experiment, cell culture medium was periodically exchanged.

### Cell lines

#### Generation of Exosc3 CKO cell lines for TC analysis

Exosc3 CKO mESCs were generated from V6.5 mESCs in our previous report.^43^ Exosc3 CKO mESCs were further edited using the CRISPR-Cas9 system to generate Exosc3 CKO/Halo-MED19 and Exosc3 CKO/Halo-MED19/mSG-Pol II mESC cell lines. Halo-tag insertion into the N-terminus of the MED19 coding region, and cell selection were performed as previously described.^24^

The resulting Exosc3 CKO/Halo-MED19 cells were further edited by inserting mStayGold (mSG)-tag into the N-terminus of the POLR2A coding region. To obtain biallelically edited cells efficiently, we generated and selected cells as detailed below: Exosc3 CKO/Halo-MED19 cells were transfected with an sgRNA and Cas9-expressing vector, along with two different donor vectors harboring PuroR-P2A-mSG or BlastR-T2A-mSG between the left and right homology arms, using the Mouse Embryonic Stem Cell Nucleofector Kit (Lonza). To increase the recombination efficiency, the donor vectors were designed to be linearized in the cells by inserting tia1l gRNA target sites at both ends of homology arms and by inserting tia1l gRNA expressing cassette in the vectors.^133^ The day after transfection, cells were treated with a low dose of M3814 (1 µM, Selleck) overnight. The culture medium was then replaced to fresh 2i containing Puromycin (InvivoGen) and Blasticidin S (FujiFilm). Successful genome-editing was confirmed by PCR using the genome extracted using QuickExtract DNA Extraction Solution (ARB-LS).

The pSNAPf-H2B sequence from a control plasmid (Addgene, # 101124) was cloned into a Piggybac vector with a G418 selection marker, followed by transposed into the endogenously tagged Halo-MED19 mESC line with G418 selection two days post Piggybac transposition.

The sgRNA and primer sequences are described in Table S1.

#### Generation of Exosc3 CKO Cell Lines for transcription imaging

To visualize bursting gene loci, we applied the MS2 strategy as previously described.^24^ Based on the Halo-MED19 tagged line, we first introduced weak ectopic MCP×2-mNeonGreen expression in cells, followed by CRISPR-mediated tagging of T2A-BSD-24×MS2 to the C-terminus of the endogenous Nanog or Sox2 loci. Cells were Blasticidin selected, single-cell sorted, and single-clone picked and expanded.

For gene-gene co-transcription imaging, we first inserted a cassette expressing MCP-mScarlet3-T2A-PCP-mSG or MCP-SNAP-T2A-PCP-mSG to Rosa 26 locus using the CRISPR/Cas9 system. The cassettes include EF1α promoter and SV40 poly(A) signal to drive MCP- and PCP-fused proteins, and SV40 promoter-driven Neomycin resistant gene terminated with HSV TK poly(A) signal. Exosc3 CKO cells were transfected with an sgRNA and Cas9-expressing vector, along with a donor vector harboring the expression cassette for MCP- and PCP-fused proteins between the left and right homology arms, using the Mouse Embryonic Stem Cell Nucleofector Kit (Lonza). The day after transfection, cells were treated with a low dose of M3814 (1 µM, Selleck) overnight. The culture medium was then replaced to fresh 2i containing G418. Next, the resulting cells were further edited by inserting T2A-BlastR-24×MS2 and T2A-PuroR-12×PP7 repeat sequences immediately downstream of the C-terminus of the coding sequence of the target genes using the CRISPR/Cas9 system: T2A-BlastR-24×MS2 to Nanog and Sox2, and T2A-PuroR-12×PP7 to Smarcad1 and Tet2. Both BlastR and PuroR contain a stop codon. We inserted T2A-BlastR-24×MS2 first, and obtained biallelically-edited cells by picking up a colony. On the next day of transfection, cells were treated with M3814 (1 µM, Selleck) for overnight and cell selection was performed using Blasticidin S (FujiFilm). In the case of Nanog editing, we performed another round of genome editing using Cas9-T2A-PuoR expression vector and an additional Nanog sgRNA vector employing [C] sgRNA technology,^134^ along with the donor vector. After the second round transfection, cells were selected using puromycin (InvivoGen). We then inserted T2A-PuroR-12×PP7 to the target genes, and selected cells with puromycin (InvivoGen).

The sgRNA and primer sequences are described in Table S1.

## METHOD DETAILS

### Imaging

#### Photoactivated localization microscopy (PALM)

The endogenously tagged Halo-MED19 was stained with 100 nM of (PA)JF549 for three hours, followed by one hour of washing in fresh media and fixation. After three washes with PBS, cells were imaged in PBS using a Nikon microscope (the same microscope described in *General setup for Single-Molecule Tracking*). The fixed-PALM configurations are as follows: 405 nm laser power = 100 mW (low-pass ND filter on, AOTF = 10%), 561 nm laser power = 370 mW (AOTF = 100%), total frames = 3000, Gain = 200, and frame rate = 20 Hz.

#### Co-localization analysis of Halo-MED19 and mSG-Pol II

Exosc3 CKO/Halo-MED19/mSG-Pol II mESC cells were plated on µ-dish 35 mm high-glass bottom dishes (iBidi) coated with PLO and laminin. For the knockout experiments, we first allow cells to settle on a coated 6 cm culture dish overnight. Cells were then washed 3 times with PBS (-), and cultured in doxycycline-free 2i media for ∼ 16 hours. Cells in the doxycyclin-free condition were plated on µ-dish 35 mm high-glass bottom dishes coated with PLO and laminin, and cultured in doxycycline-free 2i media for two nights (total doxycycline-free period is 65 hours). One day before the fixation, culture medium was replaced to fresh 2i medium containing 200 nM Janelia Fluor 646 (JF646)-HaloTag (Promega) in the presence or absence of 0.1 µg/mL doxycycline. Before fixation, cells were washed twice with 2i media without HaloTag ligand, followed by 30 min incubation in the media in the presence of Hoechst33342 (1 µg/mL, Dojindo). Cells were then fixed with 1% formaldehyde (Thermo Fisher Scientific) diluted in PBS (-) for 1 min. After quenching with 1.25 µM glycine for 5 min, cells were washed twice with PBS (-). Imaging was performed using the Nikon AXR confocal laser scanning microscope equipped with a Plan Apo Lambda 100× Oil Objective lens. After the acquisition, images were deconvolved using NIS-Elements software (Nikon). Overlaps between MED19 condensates and Pol II clusters were analyzed by creating CellProfiler pipelines^108^ (Figure S1C). Briefly, the pipeline covers identification of MED19 and Pol II clusters in each nucleus and measurement of sizes, shapes, and intensities of each cluster, and detection of overlaps between MED19 condensates and Pol II clusters. Deconvolved images obtained in 59 control cells and 40 Exosc3 KO cells were subjected to maximum intensity projection of z-stacks and then applied to the pipeline. The output tables were analyzed in R.

#### Fluorescence recovery after photobleaching (FRAP)

The endogenously expressed Halo-MED19 was stained with 100 nM Halo-JF549 (Promega, GA1111) for 20 minutes, followed by washing in fresh 2i media. We performed the FRAP experiment using ZEISS LSM980 Super-resolution Confocal (63X objective, Airyscan 2). The images were taken with 1% 560 nm excitation at 1 sec per frame. A pre-bleach frame was taken to normalize the intensity. The photobleaching power and duration were optimized to photobleach ∼95% of JF549 dye molecules. The imaging acquisition per region lasted for half a minute, ensuring the control condition recovered more than half without significant global signal loss.

#### General setup for Single-Molecule Tracking (SMT)

The SMT was performed using a Nikon Eclipse Ti microscope with a 100X oil immersion objective (NA 1.40) (Nikon, Tokyo, Japan) and an Andor iXon Ultra 897 EMCCD camera (EM gain 1,000) as previously done.^70^ The pixel size equals 160 nm on the conjugated sample plane. Before image acquisition, the cells were transferred from 2i media into prewarmed L-15 media, no phenol red (Gibco, 21083027), and were maintained at 37°C in a temperature-controlled stage during imaging. Proteins of interest were Halo-tagged and live-cell stained with Halo-(PA)JF-549 dye (a generous gift from Lavis Lab, Janelia Research Campus) for 3 hours, followed by 1 hour of washing in 2i media. For SMT, image sequences were acquired at 5 ms per frame under illumination with a minimal 405 nm for photoactivation and 561 nm (370 mW) for excitation. The 561 nm excitation power on the sample was fine-tuned by AOTF, resulting in the trajectories with around ∼10 jumps on average before photobleaching (a practical setting is AOTF = 35%). The z-position of the microscope stage was maintained during acquisition using the Perfect Focus System (PFS) of the Nikon Ti Eclipse. 5,000 to 10,000 sequential frames were taken for each region of interest (ROI).

#### Probing Mediator motion dynamics around transcriptional condensates via SMT

The endogenously expressed Halo-MED19 were co-stained with 5 nM of Halo-(PA)JF-549 (added first, 3 hours staining in total) and 100 nM of Halo-JF503 (a generous gift from Lavis Lab, Janelia Research Campus) and stain for 20 minutes (added after the (PA)JF-549 having been staining for 2 hrs 40 min). The JF503 with a 488 nm illumination was used to visualize the bulk Mediator distribution, and (PA)JF-549 was used to perform SMT of Mediator modules. For each ROI, the large, stable transcriptional condensates visible in conventional wide-field epifluorescence microscopy were focused and imaged (200 ms, 10 frames), followed by immediate SMT in the same ROI. After SMT, an additional round of bulk Mediator distribution with JF503 of the same ROI was immediately retaken for the drift correction in the downstream analysis.

#### Probing Mediator motion dynamics around bursting gene loci via SMT

Mediator molecules were labeled using 100 nM of (PA)JF549. For each ROI, five cycles of imaging were exerted: firstly, a 488 nm illumination was applied to locate and focus on a bursting Nanog locus (200 ms, 10 frames); then, SMT was performed; the Nanog locus was retaken; a moderate 405 nm (laser power = 100 mW, low-pass ND filter on, AOTF = 10%) and an intense 651 nm illumination (laser power = 370 mW, AOTF = 100%) were combined to perform live-cell photoactivated localization microscopy (PALM) with an exposure time of 50 ms for 1,200 frames; lastly, the Nanog locus was taken for the third time to confirm the bursting was still happening and provide the feasibility of drift calibration.

#### Probing dwelling dynamics of Mediator with single molecule imaging

The endogenously expressed Halo-MED19 was stained with 0.5 nM of Halo-(PA)JF-549 for 3 hours, followed by washing in fresh media for 1 hour. The same live-cell microscope system used for SMT was applied here for the dwelling dynamics measurement, except that the 561 nm laser power was tuned down to 200 mW with only 5% AOTF to reduce the photobleaching. The images were taken at 50 ms per frame without 405 photoactivation (we found that the spontaneous photoactivation was already sufficient to give us the desired visible molecule density).

#### Single-color z-stack imaging of Halo-MED19

The endogenously tagged Halo-MED19 was stained with 100 nM Halo-JF549 for 20 minutes, followed by 15 minutes of washing in fresh media and fixation with 4% paraformaldehyde (Electron Microscopy Sciences, 157-4). Cells were washed three times with PBS and imaged in PBS using the same Nikon microscope used for SMT. The microscope settings were as follows: 561 nm laser power = 370 mW, EM gain = 100, exposure time = 200 ms, AOTF percentage = 2%, z-stack range=0∼9.6 µm, and dz=0.32 µm.

#### Dual-color z-stack imaging of Halo-MED19 and H2B-SNAP

Cells were co-stained with 100 nM of SNAP-JF549 (a generous gift from Lavis Lab, Janelia Research Campus) and 100 nM of Halo-JF646 (GA1121) for 20 minutes, followed by 15 minutes of wash in fresh media and fixation with 4% paraformaldehyde (Electron Microscopy Sciences, 157-4), followed by three washes with PBS. Cells were imaged in PBS with ZEISS LSM980 Super-resolution Confocal (63X objective, Airyscan 2) with optimized pixel-size (42.5 nm) and z-stack settings (9 slices in total) and 3% laser power for both channels (gain=800).

#### Single-molecules tracking of Pol II in living cells

Exosc3 CKO/Halo-MED19/mSG-Pol II mESC cells were seeded on a glass-based dish (12 mm in diameter, 0.15 mm-thick glass; Iwaki) coated with PLO and laminin as described above. To determine the positions of Halo-MED19 labeled with HaloTag SaraFluor 650B Ligand (SF650B, Goryo Chemical) and mSG-Pol II in the nucleus, single molecules of these fluorescent probes were visualized by oblique-angle illumination on a Nikon Ti2 inverted microscope (100X 1.49 NA oil objective) equipped with dual qCMOS cameras (ORCA Quest C15550-20UP; Hamamatsu Photonics). mSG was excited using a 488 nm laser (EXLSR-488C-100-CDRH, 100 mW, Spectra-Physics) at 2.5 µW/µm^2^, and SF650B was excited using 647 nm laser (LuxX 647-140, 140 mW, Omicron) at 17.0 µW/µm^2^. Before observation, Halo-MED19 molecules were labeled with 30 nM SF650B by incubating cells for 30 min. The excitation arm included a multiple-band mirror (Di01-R405/488/561/635–25 × 36, Semrock). Emitted fluorescence signals were split into two detection arms using a dichroic mirror (Chroma: T640lpxr-UF2), with each arm fitted with a band-pass filter (ET525/50, Chroma; FF02-685/40, Semrock). The data acquisitions of dSTORM/single-molecule imaging were simultaneously performed at 37°C and at 4 ms/frame for 3,012 frames with an image size of 512 × 512 pixels. Each image was cropped to 180 × 180 pixels for analysis. The fluorescent spots were detected using the ThunderSTORM plugin of ImageJ with “Wavelet filtering” (B-Spline order = 4 and B-Spline scale = 4.0) and the “Local maximum method” (Peak intensity threshold = 30–50 and Connectivity = 8-neighbourhood). After spot detection, the post-processing steps of “Remove duplications” (Distance threshold = uncertainty) and “Drift correction” (cross-correlation with 5 bins) were further performed. The two-color images obtained from separate cameras were superimposed as previously reported.^135^ The localization precision of single-molecules of mSG and SF650B were 19.0 nm and 16.4 nm, respectively. The final pixel size was 45.5 nm.

#### Imaging of transcriptional bursts in living cells

Exosc3 CKO/Halo-MED19/MCP-SNAP/PCP-mSG/Sox2-24×MS2/Tet2-12×PP7v4 and Exosc3 CKO/MCP-mScarlet3/PCP-mSG/Nanog-24×MS2/Smarcad1-12×PP7v4 mESCs were plated on µ-Dish 35 mm high glass-bottom dishes (ibidi) coated with poly-L-ornithine (PLO) and laminin. On the day of imaging, the culture medium was replaced with fresh 2i medium supplemented with 200 nM Janelia Fluor 646 (JF646)-HaloTag ligand (Promega) and 0.1 µg/mL doxycycline (Sigma). For SNAP labeling, 60 minutes after HaloTag incubation, SNAP-Cell TMR-Star (New England Biolabs) was added at 1 µM and incubated for an additional 30 minutes. In total, HaloTag labeling was performed for 90 minutes. After labeling, cells were washed three times with fresh 2i medium and incubated in the same medium for 30 minutes. Immediately before imaging, the medium was replaced with pre-warmed L-15 containing 1 mM PD0325901 (Selleck), 3 mM CHIR99021 (Cayman), 0.1 µg/mL doxycycline, and 100 U/mL mLIF (EMD Millipore). During imaging, cells were maintained at 37 °C under humidified conditions. Imaging was performed using a Nikon AXR confocal laser scanning microscope equipped with a Plan Apo Lambda 60× oil-immersion objective lens, incubation apparatus, and a piezo stage. Images were acquired at a resolution of 512 × 512 pixels with a pinhole size of 1.2 Airy units. Z-stack images (15 slices at 0.5 μm intervals) were collected every 4 minutes for up to 4 hours. Acquired images were deconvolved using NIS-Elements software (Nikon).

### Imaging Analyses

#### Pair-correlated PALM (pcPALM)

As multiple appearances of a single protein might cause pseudo-cluster identification, we further performed pcPALM to solidify our conclusion, aiming at extracting the accurate cluster length scale in a dataset when multiple localizations from the same protein could be a potential confounding issue. Following the localization analysis (via MTT algorithm) on the Halo-MED19 fixed-cell PALM imaging series, we computed the pair-wise correlation function of the 2D localizations. The pair-correlation function of the localizations was fitted to a clustered model^136^:

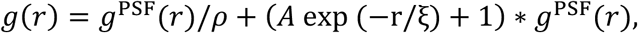

with

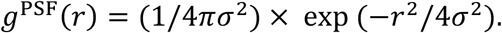

Here, *ρ* represents the average density of the protein, *A* represents the magnitude of the clustering component, *ξ* represents the length scale of the clustering component, and “∗” denotes the convolution operator. A bigger *ξ* indicates a bigger clustering size on average embedded in a dataset (Figure S1B).

#### Dynamic exchange kinetics of transcriptional condensates via FRAP

Each Halo-MED19 labeled condensate was manually cropped. The peak intensity of each condensate at each frame was obtained by fitting an isotropic 2-dimensional Gaussian function plus a background. The peak intensities of each condensate were normalized to that of the pre-bleach frame. The results from 6-10 cells in the same condition were averaged to obtain a FRAP curve (Figures 2E-H). Each FRAP curve was further fitted to an exponential equation to get the half recovery time (t_1/2_): *I*(*t*) = *A* × 2^(−*t*/*t*_1/2_ + *B*. The analysis was done with customized MATLAB code. To gain statistical insights, we performed bootstrapping with replacement on the FRAP results from 6-10 cells in each condition, and t_1/2_ was fitted in each bootstrapping. Two-tailed *t*-tests with Welch’s correction (Prism 10, GraphPad) on t_1/2_ were performed with P values as follows: DMSO (n = 6) vs DRB (n = 6), P = 0.0004 (Figure 2G); Control (n = 10) vs Exosc3 KO (n = 10), P < 0.0001 (Figure 2H).

#### SMT trajectory reconstruction and visualization

The single-molecule localization and reconstruction were done with customized MATLAB code based on the MTT algorithm.^137^ Detection settings: false-positive threshold=24, window-size 7×7pixel, and Gaussian width fitting allowed. Reconnection settings: T_off_ = 5 ms, T_cut_ = 10 ms, D_prior_ = 2 µm^2^/s, r_max_ = 270 nm. Trajectories with ambiguous connections were rejected. Trajectories with at least five jumps were kept for downstream visualization and analyses. In Figure 2L, each trajectory was randomly unicolor-coded, with the beginning labeled with a dot and the end labeled with a circle. In Figures S3A and S3B, each jump of a trajectory was color-coded by its local apparent diffusion coefficient (D = MSD/4τ), with MSD being the mean-squared displacement of each frame and τ being the time lag of each frame.

#### Mediator mobility around transcriptional condensates via SMT

For each trajectory, we performed linear regression for the MSD-τ relation and got the apparent diffusion coefficient (calculated as slope/4). We fitted each transcriptional condensate (from the bulk Mediator stain channel) with an isotopic 2-dimensional Gaussian function plus a background term, and calculated the half width at half maximum (HWHM, (2ln2)^0.5^*σ*) to define the effective range of each condensate. Based on the center positions of the condensate before and after the SMT, the trajectory positions were corrected with linear interpolation of *dx* and *dy* according to the relative time. Trajectories were kept for analysis only if the drift-corrected center of mass was within the range of HWHM to the center of the condensate. The distribution of trajectories’ logarithmic apparent diffusion coefficients can be well-fitted into a dual-Gaussian distribution (Figures 2J-K). We interpreted the peak of the lower mobility as the immobile population and the peak of the higher mobility as the diffusive population. The half maximum on the right side of the lower-mobility Gaussian and the half maximum on the left side of the high-mobility Gaussian coincided around −2.5; therefore, we cut the cumulative distribution of log *D* in two parts around −2.5, with the first part being the immobile trajectories and the second part being the diffusive trajectories (Figures 2J and 2K). Two-tailed *t*-tests with Welch’s correction (Prism 10, GraphPad) were performed with p values as follows. Bound: DMSO (n = 531) vs DRB (n = 472), P = 0.9168 (Figure 2J). Diffusive: DMSO (n = 315) vs DRB (n = 257), P = 0.0068 (Figure 2J). Bound: Control (n = 120) vs Exosc3 KO (n = 120), P = 0.8844 (Figure 2K). Diffusive: Control (n = 55) vs Exosc3 KO (n = 67), P = 0.018 (Figure 2K).

#### Mediator exchange dynamics of transcriptional condensates via SMT

We fitted each transcriptional condensate (from the bulk Mediator stain channel) with an isotopic 2-dimensional Gaussian function, and the fitted *σ* defined the condensate core (∼250 nm). In each condition, we identified trajectories of three types: entering, exiting, and confined. For “entering” trajectories, the first localization is outside of the condensate core, and the last localization is inside of the condensate core; for “exiting” trajectories, the first localization is inside of the condensate core, and the last localization is outside of the condensate core; for “confined” trajectories, all localizations are inside of the condensate core (Figure 2L). We then used the “confined” trajectories as the baseline and calculated the relative enrichment of traveling trajectories, which is defined as the ratio between “entering” and “confined” trajectories or the ratio between the “exiting” and “confined” trajectories (Figure 2M). The standard deviation of such relative enrichment values was further estimated by assuming a binomial distribution (Figure 2M). To gain statistical insights, we performed Chi-Square Test by creating a 2×2 contingency table with p values computed as follows: for “entering” trajectories, DMSO (n = 486) vs DRB (n =4 30), P = 0.04; for “exiting” trajectories, DMSO (n = 482) vs DRB (n = 431), P = 0.18; for “entering” trajectories, Control (n = 110) vs Exosc3 KO (n = 140), P = 0.02; for “exiting” trajectories, Control (n = 110) vs Exosc3 KO (n = 140), P = 0.07.

#### Dwell time of Mediator single molecules in nuclei

Following the localization analysis (via MTT algorithm) on the Halo-MED19 single molecule imaging time series, we performed the DBSCAN with a short neighbor connection threshold (r_max_ = 60 nm) and high minimum neighbor numbers (n_min_ = 80) to recognize clusters that potentially contain immobile and long-lived molecules. The recognized clusters were taken out from the original localization map, and the remaining localizations underwent several deflation loops of DBSCAN with declining minimum neighbor numbers (n_min_ = 40, 20, 10, 5, 3), where more and more clusters were recognized and taken out from the localization map in each loop. After this spatial clustering, we further performed temporal clustering on each spatial cluster, where the time-correlated single-molecule events were extracted: if no more than two dark frames temporarily separated two consecutive localizations, they would be grouped into the single-molecule event. We further filtered out the marginal single-molecule events that appeared in the first frame or lasted until the last frame of an ROI. The duration of each single-molecule event is counted, and the cumulative probability of durations of all single-molecule events from the same condition was fitted into a dual-exponential function 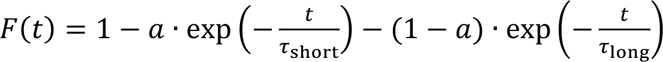, where two timescales were fitted. We focused on the apparent dwell time of the long timescale (τ_long_) of transcriptional proteins as previous work did,^70^ where the calibrated dwell time (τ_long,cali_) of a live-cell sample was calculated from a fixed sample: 1/τ_long,cali_ = 1/τ_long,live_-1/τ_long,fix_. We performed downsampling with a 0.2 sampling ratio thirty times, and the calibrated dwell times of the sample condition were pooled together for performing *t*-tests with boxplots and p values reported (Figures S3D-E).

#### Boundary sharpness of condensates

The maximum intensity projection (MIP) was created from the single-color z-stack imaging of Halo-MED19. The peaks in each MIP were identified with a spline wavelet filter.^138^ Each peak is fitted with an isotropic 2D Gaussian function with an additional background term, and the boundary sharpness is defined as the peak intensity (arbitrary unit) divided by the fitted full width at half maximum (FWHM, 2(2ln2)^0.5^*σ*) (Figure 3F). Two-tailed *t*-tests with Welch’s correction (Prism 10, GraphPad) were performed to evaluate the statistical significance.

#### Chromatin accessibility of condensates

We selected cells with comparable JF549-H2B intensity for imaging and analysis, and the brightnesses in different z-stacks were further normalized according to each z-stack’s bright pixels (mean of pixels in between the [0.85,0.99] quantiles). This normalizing operation was done for both channels. For each z-stack, we computed the MIP of the JF646-MED19 channel. Based on the MIP, we performed the peak identification (with the spline wavelet filter) to find the center of each peak in the x-y plane (x_0_,y_0_), from which we further scanned along the z direction in the z-stack to locate the z position where the MED19 signal peaked (z_0_). Centered at (x_0_,y_0_ z_0_), we made a 2D cropping in the x-y plane for both channels. In this dual-channel 2D cropping, the MED19 signal was used to fit an isotropic 2D Gaussian function with an additional background term, and the full width at half maximum (FWHM, 2(2ln2)^0.5^*σ*) was calculated to define a local zone (x-x_0_)^2^+ (y-y_0_)^2^< FWHM^2^. This local zone included both the Mediator condensate and the surrounding background. The pixel-wise brightness of the JF646-MED19 channel was partitioned into eight levels, and the JF549-H2B brightnesses of the same pixels were extracted (Figures 3I-J). These results include such pixel-wise analysis pooled from local zones of different condensates from more than twenty cells in each condition, with the mean and standard error of the mean (SEM) plotted. Our analysis suggested that in both the transcription inhibition and the RNA exosome knockout conditions, the H2B intensity decreases (or is absent) when the pixel-wise MED19 intensity goes high.

#### Area quantification of super-resolved clusters

Following the localization analysis (via MTT algorithm) on the Halo-MED19 fixed-cell PALM, we performed DBSCAN with the neighbor connection threshold r_max_ = 105 nm and declining minimum neighbor numbers (n_min_ = 12, 11, 10, …, 5, 4) in a multiple deflation-loop procedure as mentioned in the subsection *Dwell time of Mediator single molecules in nuclei*. More and more valid spatial clusters were extracted throughout multiple deflation loops of DBSCAN. A convex polygon was created based on the localizations of each cluster, whose area was used to infer the size of super-resolved Mediator clusters (Figure 1B).

#### Colocalization analysis between MED19 clusters in dSTORM movies and single Pol II molecules

The distances between the centroid of Pol II molecules and the contour of MED19 clusters were quantified using single-molecule tracking data of the Pol II and binarized images of MED19 clusters in dSTORM movies. To generate binarized images of MED19 clusters in the dSTORM movies, we used the kernel density estimation (KDE) method as previously reported.^139^ This analysis was performed using colocalization analysis software developed by our group.^140^

### Phase-Field Simulations

#### Two-component reaction-diffusion model of transcriptional protein and RNA

We constructed a phenomenological free energy functional following the previous Landau model framework.^37^ Our model assumes that the transcriptional protein concentration field *φ*_p_(***r***,*t*) self-attracts at intermediate concentrations (favoring condensation) and self-repulses at high concentrations (volumetric exclusion). The RNA concentration field *φ*_r_(***r***,*t*) self-repulses at high concentrations. There is an attractive interaction between *φ*_p_ and *φ*_r_ at low concentrations and a repulsive interaction at high concentrations. There is also a penalty of imbalanced charge between *φ*_p_ and *φ*_r_. Lastly, protein also has a gradient energy penalty reflecting the additional protein interaction on the surface of a transcriptional condensate. We define the free energy functional (normalized to *k_B_T*= 1) as:

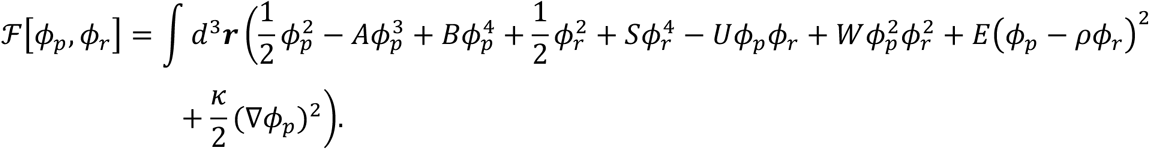

The integral is taken over the whole 3-dimensional simulation space. *A*, *B*, *S*, *U*, *W*, *E*, and *κ* are parameters that determine the strength of different interactions. *ρ* is the relative charge of RNA to protein. The terms ½*φ*_p_^2^ and ½*φ*_r_^2^ ensure that protein and RNA molecules obey the basic Fick’s laws of diffusion when there are no additional interaction terms. Therefore, the free energy functional can be rewritten as

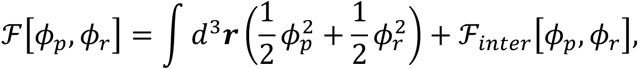

where:

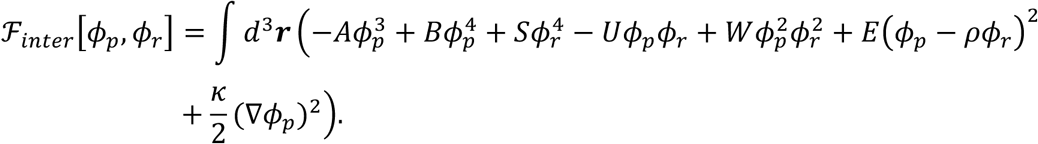

The chemical potentials of the protein and RNA components owing to the additional interaction terms are calculated as follows:

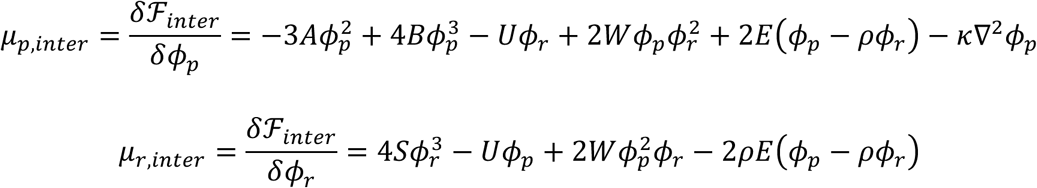

It has been reported that interactions between transcriptional proteins and RNA may generally exist.^37,70^ Here, we model such general protein-RNA interactions as transient complexes (*φ*_pr_), which follow a first-order reaction at equilibrium at each local position ***r*** (or, say, voxel, in a simulation space):

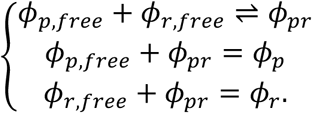

Given a specific disassociation constant *K_d_* and a pair of *φ*_p_ and *φ*_r_ of a voxel, the I_)*_ of the voxel can be solved analytically:

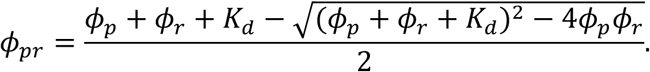

Accordingly, the non-complex concentrations of protein and RNA are ϕ*_p_*_,*free*_ = ϕ*_p_* - ϕ*_pr_* and ϕ*_r_*_,*free*_ = ϕ*_r_* - ϕ*_pr_*, and the non-complex fractions of protein and RNA can be calculated as *f_p_*_,*free*_ = ϕ*_p_*_,*free*_/ϕ*_p_* and *f_r_*_,*free*_ = ϕ*_r_*_,*free*_/I*_r_*, respectively. Thus, the chemical potential of the transient protein-RNA complexes owing to the additional interaction terms can be expressed as:

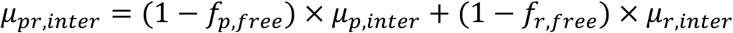

The reaction-diffusion dynamics of protein and RNA are:

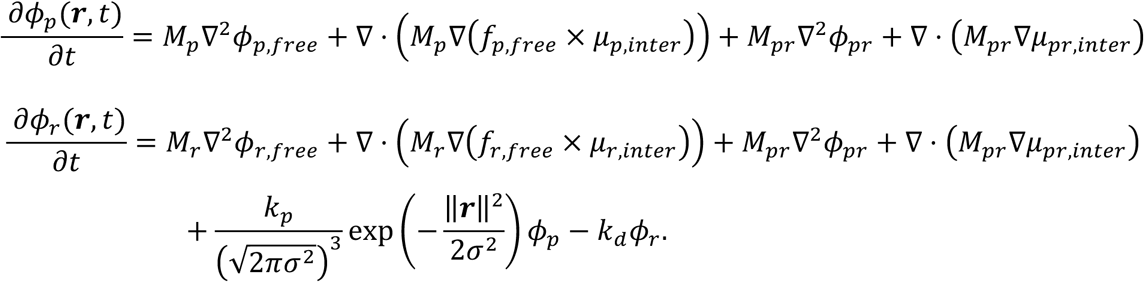

*M_p_*, *M_r_* and *M_pr_* are the mobilities of protein, RNA, and transient protein-RNA complexes, respectively. *k_p_* is the RNA production rate, and *k_d_* is the RNA degradation rate. We further assume the transcription only occurs around the origin (e.g., an active promoter at ***r*_0_**=(0,0,0)) and model this transcribe region as a three-dimensional Gaussian function with an isotropic spatial extent of *σ* ^141^ (in the actual simulation, we used a truncated Gaussian at half maximum, meaning the value drop to zero when ||***r***||>1.18*σ*). The product of the promoter and protein concentrations determines the RNA transcription activity.

#### Simulation parameter settings

Unless otherwise mentioned, the simulations were carried out with the following parameters (i.e., “Control” condition). Simulation space: L_x_ = L_y_ = L_z_ = 31 units length. Voxel dimension: (1 unit length)^3^. Initial condition: 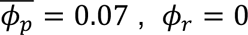 everywhere. Boundary conditions: no-flux boundary for protein (*φ*_p_) and absorbing boundary for RNA (*φ*_r_). Time increment per step: dt = 1.25E-4. Interaction parameters: *A* = 10, *B* = 5, *S* = 15, *U* = 5, *W* = 1, *E* =1.5, *κ* = 30, *ρ* = 1, and *K_d_* = 1. Mobilities: *M_p_* = 1, *M_r_* = 0.3, and *M_pr_* = (*M_p_*M_r_*)/(*M_p_*+*M_r_*) (harmonic mean). Reaction terms: *k_p_* = 2,000, *σ* = 2, and *k_d_* = 1. The simulations are done with customized code in MATLAB (R2024b, MathWorks).

We further explored several scenarios by changing some parameters from the Control condition. For the “synthesis OFF” condition, we set *k_p_* = 0, and other parameters remain unchanged. For the “Exosome KO” condition, which was reported to lead to both reduced RNA degradation and prolonged retention,^39,43,142^ we decreased both *k_d_* and *M_r_* by five folds, *k_d_* = 0.2 and *M_r_* = 0.06, while other parameters remained unchanged. We further investigated how active synthesis, degradation, and mobility of RNA can alter the physical properties of a transcriptional condensate: starting from the “Control” condition, we also performed three single-variable scans along three axes, *k_p_*, *k_d_*, or *M_r_* (Figures S2B-D). Across the range of RNA production (*k_p_*), which shows non-monotonic changes in condensate intensity (between “synthesis OFF” and “Control”), almost monotonic decrease in incorporate rate and intra-condensate exchange were observed. Decrease in RNA production (*k_d_*) from the “Control” condition accompanied decrease in intensity, incorporate rate, and intra-condensate exchange. As for RNA mobility (*M_r_*), intra-condensate exchange was sensitive to changes in RNA mobility, whereas the effects on intensity and incorporate rate were modest.

#### Condensate intensity

For the average intensity of proteins in a steady-state condensate, we first defined the condensate boundary, which is a sphere with intensity of 0.05 (the core of the condensate has an average intensity of about 1.0 to 1.2). We then calculated the voxel-wise average intensity of the condensate, followed by normalizing it to the synthesis OFF case (Figure S2B).

#### Protein incorporation dynamics into a condensate

Similar *in silico* “isotope tracing” was performed to investigate the protein incorporation dynamics into condensate with different RNA kinetics. After the system reached a steady state, we labeled the existing protein field as isotope 2 (i.e., 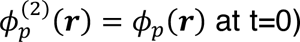. We then added a tiny amount of protein isotope 1 in the dilute phase outside the periphery of the condensate:

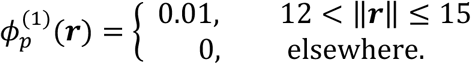

The average intensity of protein isotope 1 inside of the condensate (‖***r***‖ ≤ 9) was recorded every 0.002 simulation time. This intensity series from each simulation was normalized to [0,1] according to its minimum and maximum values. The inverse of the half-turnover time of the protein isotope 1 was reported as an estimator rate of the protein incorporation dynamics into a condensate (Figures 2D and S2C, S2E).

#### Protein intra-exchange dynamics in a condensate

To study the protein motion in condensate, we performed “isotope tracing” *in silico*. We first let the simulation run into a steady state. We then digitally “labeled” the protein field around the core of the condensate (‖***r***‖ ≤ 4) as isotope 1, and the protein field elsewhere as isotope 2. Mathematically, the initial distributions of these two protein isotopes are:

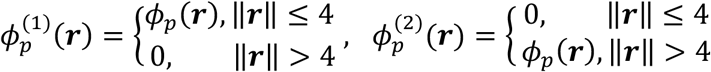

Both isotopes follow the same diffusion dynamics to evolve:

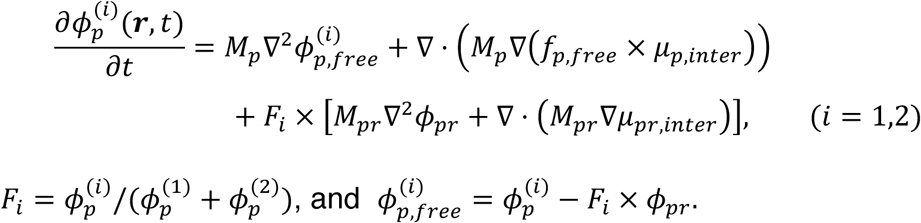

The reaction-diffusion dynamics for the RNA field remain unchanged and only depend on the sum of both protein isotopes rather than individual isotopes. The average intensity of protein isotope 1 near the center (‖***r***‖ ≤ 3.5) was recorded every 0.02 simulation time. This intensity series from each simulation was normalized to [0,1] according to its minimum and maximum values. The inverse of the half-turnover time of the protein isotope 1 was reported as an estimator rate of the protein intra-exchange dynamics in a condensate (Figures 2D and S2D, S2F).

#### Boundary sharpness

For the boundary sharpness, we defined it as the peak intensity divided by the full width at half maximum (FWHM). Such values are further normalized to the synthesis OFF case. The total RNA density inside of a condensate varies according to the RNA synthesis rate *k_p_*, and we found that for any given comparable total RNA density inside of the condensate, the RNA exosome KO case—represented by decreased degradation rate and mobility of RNA—always led to higher condensate boundary sharpness compared with the case with normal RNA exosome functionality (Figure S3H). We picked the RNA exosome KO case where the total RNA density inside of the condensate matches that of the control condition for illustration (Figure 3A).

#### Chromatin permeability

We also wonder how the material properties of the condensate can be altered by RNA kinetics. We adopted the mathematical description of the mechanical exclusion of chromatin by droplet formation from the previous field simulation work.^143^ The interaction part of the new free energy functional is slightly modified as:

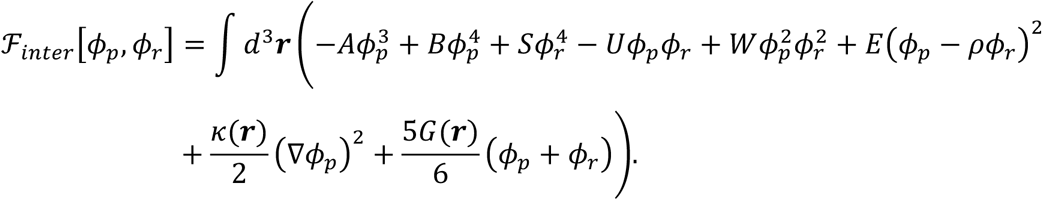

Here, G(***r***) is an incompressible elastic medium of Young’s modulus representing chromatin. *k*(***r***) = 2(λ_0_ + 2*r*_mesh_*G*(***r***)/9)^$^ is the new surface penalty term modulated by s(M). With those modifications, the chemical potential of the interaction terms now becomes:

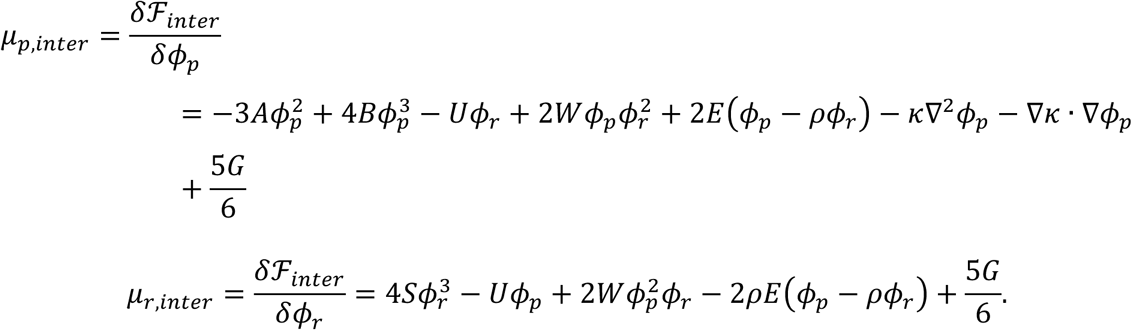

The parameter setting in our simulation is λ_0_ = √5 and ***r***_mesh_ = 7.5. As for G(***r***), we set it as a sphere with a blurred surface:

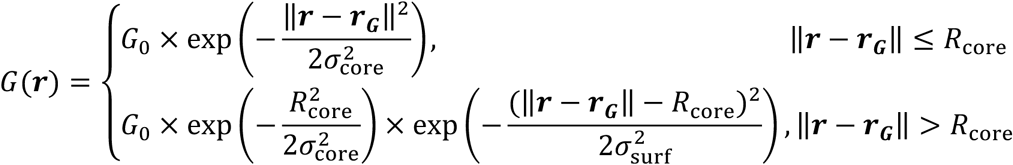

where *G*_0_ = 1, *R*_core_ = 8, *σ*_core_ = 20, and *σ*_surf_ = 1. To avoid the permeability measurement being derealized by the immobilized transcriptional locus in our simulation, we simplified the RNA synthesis term by assuming the transcription can occur wherever the protein density field exists ^37^; thus, the reaction part in the reaction-diffusion equation of RNA becomes: *k_p_ϕ_p_* - *k_d_ϕ_r_*. Accordingly, the Control and Exosome KO conditions now have an RNA synthesis rate *k_p_* = 2, and the synthesis OFF condition still has *k_p_* = 0. To further simplify the time and length scales involved in this simulation, we assumed equal mobility for protein and RNA in the Control condition with *M_p_* = *M_r_* = 1, and the Exosome KO condition has a five-fold lower RNA mobility, *M_r_* = 0.2.

We first prepared a steady-state condensate in the default simulation space (L_x_ = L_y_ = L_z_ = 31). At the beginning of the *in-silico* experiment, we placed the prepared condensate on the left side of a rectangular cuboid (L_x_ = 61, L_y_ = 31, L_z_ = 31), and placed the center of the chromatin sphere (***r_G_***) at the right side of the rectangular cuboid. As the simulation continued, the chromatin sphere gradually moved to the left and intruded the condensate, and the total protein quantity penetrated into the chromatin sphere was recorded over time. The inverse of the half-penetration time was reported as the chromatin penetrating rate (Figures 3B-C).

### Growth curve assay

Exosc3 CKO/Halo-MED19/mSG-Pol II cells were seeded at 5 × 10⁴ cells (KO and Reverse) or 1.5 × 10⁵ cells (Ctrl) in 2i medium. Twelve hours after seeding, cells were washed three times with PBS (-). The cell number at this point was measured and defined as time 0. Washed cells were then cultured in either doxycycline-free (KO) or doxycycline-containing (Ctrl) 2i medium, and cell growth was measured at 24, 48, and 60 hours. After 60 hours in doxycycline-free medium, the culture medium was changed to either doxycycline-free (KO, continued) or doxycycline-containing (Reverse) 2i medium, and growth was measured again at 24, 48, and 60 hours. Fold growth was calculated relative to the cell number at time 0.

### Library preparation for multi-omics analyses

#### ChIP-seq library preparation

ChIP-seq was performed as previously described^144^ (available at http://younglab.wi.mit.edu/hESRegulation/Young_Protocol.doc), with minor modifications. We followed the protocol unless otherwise described specifically below. Total of 1 ×10^8^ cells per immunoprecipitation were cross-linked with 1% formaldehyde for 10 minutes at room temperature. Cells were then lysed, and the nuclei were resuspended in SDS-shearing buffer (10 mM Tris-HCl pH 8.0, 1 mM EDTA, 0.15% SDS, 1 x cOmplete protease inhibitor cocktail). Chromatin was sonicated using the Bioruptor II (BMBio, BR2012A TYPE12) for 30 min (30 seconds on, 30 seconds off; high power load) to obtain an average fragment size of 300–500 bp. Triton X-100 was then added to a final concentration of 1%, and the sample was centrifuged at 20,000 × g for 10 min at 4 °C. Chromatin immunoprecipitation was then performed for overnight at 4 °C using 10 µg anti-Pol II (BioLegend), 10 µg anti-MED1 (Bethyl), 5 µg anti-DSIF (BD Biosciences) or 1.37 µg anti-NELF (Cell Signaling) antibodies pre-conjugated to Protein G Dynabeads (Thermo Fisher Scientific). After five washes in Wash buffer and following one wash in TE/ 50mM NaCl buffer, immunoprecipitated DNA was treated with RNase A (Thermo Fisher Scientific) and eluted while reverse-crosslinking in the presence of proteinase K (Thermo Fisher Scientific). Eluted DNA was purified using the QIAquick PCR Purification Kit (QIAGEN). ChIP-seq libraries were prepared using the TruSeq ChIP Sample Prep Kit-Set A (Illumina) with modifications. Briefly, ChIP DNA was end-repaired, purified using AMPure XP beads (Beckman Coulter), and eluted. A-tailing was performed, followed by adapter ligation. Double size selection was conducted to remove large fragments and adapter dimers. Library fragments were amplified by PCR, purified again, and sequenced on an Illumina HiSeq X platform (Macrogen Japan Corp.) with paired-end 150 bp reads.

#### PRO-seq library preparation

PRO-seq was performed according to the published PRO-seq protocol.^145^ Briefly, isolated nuclei from Exosc3 CKO cells cultured in the presence or absence of 0.1 µg/mL doxycycline were permeabilized and subjected to nuclear run-on in the presence of biotin-11-CTP (Perkin Elmer). After the run-on, RNA was extracted and fragmented, and biotinylated RNAs were enriched using Dynabeads Streptavidin M280 beads (Thermo Fisher Scientific). The enriched RNAs were ligated with 3′ and 5′ adaptors, and then reverse-transcribed. After PCR, libraries were size-selected by gel extraction. The libraries were sequenced on Illumina HiSeq 2500 PE100 platform by Rarevariant Inc.

#### RNA-seq library preparation

Exosc3 CKO mESCs were directly lysed in TRIzol (Thermo Fisher Scientific), and RNA isolation was performed using the Direct-zol RNA Microprep Kit (Zymo Research) following the manufacturer’s instructions. Total RNA was used for constructing stranded RNA libraries with the rRNA Depletion Kit (MGI) and RNA Directional Library Prep Set (MGI) according to the manufacturer’s protocols. The libraries were sequenced on an MGI DNBSEQ-G400 platform (Rarevariant Inc) with paired-end 150 bp reads.

#### scRNA-seq and snRNA-seq library preparation

scRNA-seq and snRNA-seq library preparation was performed using Chromium Next GEM Single Cell 3′ Reagent Kits v3.1 or Chromium Next GEM Single Cell 5′ Reagent Kits v2 (10X Genomics), respectively, according to the manufacturer’s protocol (CG000315 RevE and CG000331 Rev E, respectively). For 5′ analysis, nuclei isolation was performed according to the 10X Genomics’s protocol (CG000124 Rev F). In both experiments, debris and cell/nuclei clumps were removed by passing through 40 µm Flowmi Tip Strainer (SP Bel-Art). The libraries were sequenced on Illumina NovaSeq 6000 PE150 platform by TaKaRa. Fastq files were generated using bcl2fastq and cellranger mkfastq.

#### Hi-C library preparation

Hi-C library preparation was performed following the iconHi-C protocol ver. 1.1^146^ (available at https://figshare.com/articles/online_resource/iconHi-C_protocol_v1_1_pdf/14669751) developed by the Laboratory for Phyloinformatics, RIKEN BDR with minor modifications. We followed the protocol unless otherwise described specifically below. Briefly, Exosc3 CKO cells were pelleted and immediately fixed in 1% formaldehyde and 2 mM DSG for 10 min at 25°C, followed by quenching with 1/20 volume of 2.5 M glycine. Cells were washed twice with ice-cold PBS and stored at −80°C until further processing. Fixed cells were permeabilized with NP-40-containing buffers and digested with DpnII and HinfI at 37°C for ∼16 h with gentle shaking. Biotinylated nucleotides were incorporated at the digested DNA ends using Klenow DNA polymerase, followed by proximity ligation with T4 DNA ligase for 4-6 h at 16°C. The ligated DNA was purified by phenol-chloroform extraction and ethanol precipitation. RNA contamination was removed by RNase A treatment, and proteins were digested using Proteinase K. Hi-C libraries were prepared through fragmentation using a Covaris sonicator, size selection with AMPure XP beads, and enrichment of biotinylated DNA using streptavidin-coated beads. End-repair, A-tailing, and adapter ligation were performed using the KAPA LTP Library Preparation Kit, followed by PCR amplification. Sequencing was performed on the HiSeqX platform (Illumina) and 528-603 million read-pairs were obtained.

### Bioinformatics analyses for multi-omics data

#### ChIP-seq data processing

All computational analyses were carried out using UCSC (GRCm38/mm10) mouse gene annotations. The raw reads were trimmed using fastp with options −q 15 −n 10 −u 40. The trimmed reads were aligned to mm10 using bowtie2 by the end-to-end mode. Duplicated reads were filtered using the Picard MarkDuplicates function. Bigwig files were generated using deepTools^113^ bamCoverage with options −-binSize 50 −-normalizeUsing RPGC −-effectiveGenomeSize 2150570000 −-ignoreForNormalization chrX –extendReads −bl mm10-blacklist.v2.bed (a bed file listing the ENCODE blacklist regions, obtained from https://github.com/Boyle-Lab/Blacklist). The signal distribution in the regions of interest was computed using deepTools computeMatrix or bedtools muticov. We also utilized H3K27ac and H3K9me3 ChIP-seq data in 2i-cultured mESCs from *Yang et al.*, 2019.^107^

#### ChIP-seq peak and super-enhancer calling

Before peak calling, all replicates of MED1 ChIP-seq and input data were merged using samtools merge. The merged BAM files were then processed with macs3 using the options −f BAMPE −g mm −q 0.01. BAM files of genomic input controls were also provided to macs3 using the −c option. Peaks detected in the ENCODE blacklist regions (mm10-blacklist.v2.bed, obtained from https://github.com/Boyle-Lab/Blacklist) were removed using bedtools intersect. Super-enhancers were subsequently called using the merged BAM files using ROSE algorism^6,29^ with the options −s 12500 −t 2500. The gene closest to a given SE is defined as an SE-associated gene.

#### PRO-seq data processing

After trimming adaptors using cutadapt^114^ requiring a minimum read length of 10 bp, reads derived from rRNA were filtered using bowtie2.^109^ The remaining reads were aligned to mm10 using bowtie 2 with options −-very-sensitive −X 350. The 5′ ends of aligned secondary reads in proper pair (3′ end of RNA) were used for coverage analysis using bedtools^113^ genomecov. The resulting 3′ end count files are used to generate bigwig files using bedGraphToBigWig. BigWig files were also generated from the paired-end read fragment using deepTools^113^ bamCoverage with the filterRNAstrand option, RPKM normalization, and bin size of 5.

#### RNA-seq data processing

Paired end reads were first mapped to ribosomal RNA and various repetitive sequences such as U1 snRNA using Bowtie2,^109^ and then subsequently mapped to the mouse mm10 transcriptome and genome using STAR aligner.^110^ The ensuing reads were filtered for uniquely mapping and properly paired reads. BigWig files were also generated using deepTools^113^ bamCoverage with the filterRNAstrand option, RPKM normalization, and bin size of 5.

#### de novo Transcriptome Assembly

The RNA transcriptome was assembled *de novo* using the Stringtie algorithm,^117^ followed by various filtering steps to categorize transcript classes (Figure S4D). For instance, uaRNAs were defined as divergent transcripts with a 5′-prime end within 1 kb upstream and antisense of the closest gene TSS.^147^ Enhancer RNAs (eRNAs) were defined as transcripts overlapping a 1 kb window of an MED1 enhancer region (super-enhancer or typical enhancer).

Genomewide identification of non-coding RNAs was performed as previously described, ^43^ with minor modifications. Stringtie^117^ was run on each RNA-seq dataset using the parameters –f 0.1 –c 5 –g 10 –j 15 and mm10 protein-coding genes as the reference annotation. The resulting candidate transcripts in either Watson or Crick orientation were separately merged into two non-redundant sets of transcripts using the merge mode of Stringtie with the parameters –F 0.1 and –T 0.1 and mm10 protein-coding genes as the reference annotation, and further merged into one non-redundant sets of transcripts. These transcripts were first removed for any transcript that overlapped mm10 genes, snoRNAs, and known miRNA genes. Any candidate transcripts were then aligned against the antisense version of mm10 genes, and majority of them was categorized into antisense RNAs: antisense transcripts that did not overlap the TSS. After filtering convergent RNAs: those that started within the gene and was transcribed across the TSS, the remaining candidate transcripts were further analyzed for uaRNAs: transcripts that were antisense to the coding gene and started within 1 kb of the TSS. The remaining candidate transcripts were further subsegmented into super-ehancer RNA (seRNA) or typical enhancer RNA (teRNA): transcripts that overlapped a flanking 1 kb window of super-enhancers or typical enhancer identified in MED1 ChIP-seq analysis. Finally, the remaining candidate RNAs were filtered for *de novo* lncRNAs by removing previously annotated lncRNAs followed by running the Slncky algorithm.^148^ We identified 684 high-confidence uaRNAs. While this number is less than the number of uaRNA in FBS condition analyzed in our previous paper,^43^ this may be partly explained by the increased pausing levels and reduced Myc levels in 2i conditions and the effects of serum.^149^

#### Differential Expression Analysis of PRO-seq and RNA-seq

The number of PRO-seq and RNA-seq reads per transcript was counted by using HTSeq (v2.0.9)^150^ in intersection-strict mode. After filtering out the genes with a maximum count per million (CPM) across all samples of less than 0.1, differential transcripts were called using edgeR, where we normalized libraries using UQ normalization.

#### Metaplots

We filtered the intervals for metaplot as follows. For metaplots at enhancers, we aligned against centers of MED1-defined enhancer summits of SE and TE constituents. Any overlapping enhancer summits within a 2.5 kb window of TSS of genes were excluded for ChIP-seq metaplots. To avoid interference from nascent signals of uaRNAs and pre-mRNAs, and eRNAs produced from closely distributed enhancers, we filtered out any overlapping enhancer summits within a 3 kb window and also any that overlapped a TSS – 2.5 kb ∼ TES region of genes. Enhancers with poor evidence of eRNA production were further filtered by referring *de novo* transcriptome assembly data. For metaplots around TSS, we first identified isoforms with the TSS having the strongest PRO-seq peak using Pausing_Index.py.^52,151^ The minimum required PRO-seq TSS peak intensity (RPM) was set to 0.01. Genes shorter than 10 kb were excluded, and any overlapping genes were removed.

To create metaplots for ChIP-seq, RPGC (reads per genome coverage)-normalized and input-subtracted bigwig coverage files were generated using deepTools bamCoverage. These files were applied to deepTools computeMatrix with a bin size of 100. PRO-seq metaplots were created similarly, but paired-end read fragment coverage files were generated using deepTools bamCoverage with the filterRNAstrand option and RPKM normalization. deepTools computeMatrix was then performed with a bin size of 20.

#### Pausing Indices

Bed files used to calculate the pausing index were generated from a gtf file of protein-coding genes using a modified script originated from extract_genomic_locations.py,^52,151^ and the regions shorter than 2500 bp were removed. Bedtools map was utilized to obtain strand-matched PRO-seq count in the regions of interest using the bedtools genomecov results as an input. To calculate pausing index, we made modifications to Pausing_Index.py^52,151^ to enable operation without input results and compatibility with the bedtools map count file format. The definition of pausing index is: signals around the strongest TSS (TSS ∼ TSS + 100 bp) divided by gene body signal (TSS + 250 bp ∼ TSS + 2250 bp).

#### 3′-scRNA-seq data processing and expression variance analysis

Reads were mapped to mm10 and parsed to determine the sample barcode associated with each cell using CellRanger v7.1.0.^127^ The resulting matrices were processed using Seurat v5^128^ by generating Seurat Objects with min.cells of 3 and min.features of 50. We excluded cells with the percentage of mitochondrial UMI is greater than 10 and the number of features is less than 3000. Counts were log-normalized, and the variable features were determined using FindVariableFeatures function. After running ScaleData, and RunPCA, data integration was performed using Harmony, and the first 45 PCs were used to run FindNeighbors. Clusters were identified using FindClusters with a resolution parameter of 0.18. Cellular differentiation status was defined based on the expression of sets of pluripotency markers (Nanog, Klf4, Esrrb, Pou5f1, Sox2, Dppa5a, and Zfp42), primed markers (Fgf5, Otx2, Car4), and differentiated markers (epiblast: Krt8 and S100a6; primitive endoderm: Col4a1, Gata4, and Gata6; mesoderm: Hand1, T, Twist2, Foxa2, and Mixl1). Estimation of the biological variance of gene expression among cells was performed using the scran^129^ modelGeneVar function, which decompose total variance into biological and technical components. In Figures S5B and S5C, genes were classified into three categories —Low (0.3 ∼ 2.5), Middle (2.5 ∼ 4), and High (> 4)—based on their expression levels.

#### 5′-snRNA-seq data processing

CellRanger’s default genes.gtf for mm10 (refdata-gex-mm10-2020-A) was merged with a gtf file of the *de novo* transcriptome assembly results using cellranger mkref. This file was specified as a transcriptome when mapping reads to mm10 and parsing the sample barcode associated with each cell using CellRanger v7.1.0.^127^ The resulting matrices were processed using Seurat v5^128^ by generating Seurat Objects with min.cells of 3 and min.features of 50, and then excluded cells with the percentage of mitochondrial UMI is greater than 10.

#### eRNA-gene co-expression analysis

To account for multiple eRNAs derived from distinct SE regions but associated with a single gene, eRNAs were aggregated per gene. The integration was performed by summing the expression values across all seRNAs associated with the gene. The resulting matrices were then binarized by setting counts other than 0 to 1. For each gene-seRNA pair, the Pearson correlation coefficient was calculated, and the P values were calculated using the chi-square test. The Benjamini–Hochberg correction was applied to the P values to obtain false discovery rate. Significant co-expression events were determined based on the criteria FDR < 0.05 and Pearson correlation coefficient > 0.1. These processes were performed by adapting the pipeline used in the previous study.^59^ The proportion of cells where both the gene and seRNA were expressed was determined. The proportion of seRNA-expressing cells among gene-expressing cells were also calculated.

#### Hi-C data processing

The raw reads were trimmed using Trim Galore with options −e 0.1 −-phred33 −q 30. The trimmed reads were aligned to mm10, and Hi–C contact matrices and valid pair files were generated using HiC-Pro^118^ with various bin sizes. Conversion of matrix files and valid pair files formats were conducted using HiCExplorer^119^ and HiC-Pro utilities. PC1 was calculated using HiC1Dmetrics.^120^ Differential interchromosomal interactions were analyzed using Homer analyzeHiC with an option −res 20000. TAD calling, read count normalization, and KR correction were conducted using HiCExplorer. For contact data visualization, CoolBox,^123^ karyoplotR,^124^ and circlize^125^ were used. Compartment networks were drawn using igraph.^126^

#### Compartment classification and analysis

Compartments were classified into three categories—A, B, and undefined—based on PC1 values. A hierarchical annotation approach was applied to further classify compartments. First, compartments containing SEs were defined as SE compartments. Among the remaining compartments, those containing TE were classified as TE compartments, while the rest were categorized as undefined compartments. Pairwise changes in interactions between compartments on the same chromosome (Exosc3 KO vs Ctrl) were computed using observed-over-expected values at a 200 kb resolution, where observed values were read count-normalized and KR-corrected, and expected values were calculated as the mean contact frequency for all positions at the same genomic distance. Compartments smaller than the resolution limit (< 200 kb) were excluded. Pairwise contact change scores were obtained for each compartment, and mean values were calculated. Each compartment was classified as “Up” or “Down” based on intra-chromosomal interaction changes: compartments with a down ratio greater than 0.5 were labeled “Down”, while those with an up ratio greater than 0.5 were labeled “Up”. The ratio and bias were calculated as:

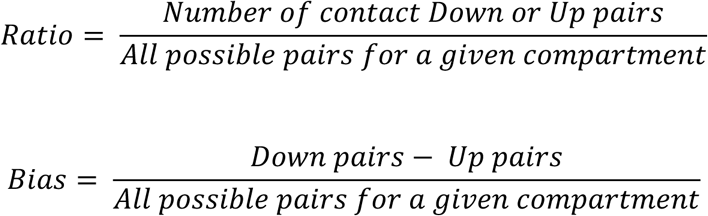

For pairwise analysis of inter-chromosomal interactions, interactions with FDR < 0.05 were considered significant. Fractions of SE-SE and TE-TE compartment pairs calculated for all possible combinations (24,143 and 3,194,372 pairs, respectively) and observed significant interactions (21 and 257 pairs, respectively). Links for all SE-SE pairs as well as SE-SE pairs with changes more than 0.2 compared to control were visualized using circlize package^125^ in R.

Links between SE-containing A (SE-A) compartment pairs exhibiting decreased contacts (changes < −0.2) were visualized using karyoplotR package^124^ in R.

#### Gene-gene correlation analysis

We removed mitochondrial mRNAs from count matrices, and the matrices were processed using scprep as follows: after filtering cells by library size (cutoff = 500), counts were normalized by library size and then applied to square root transformation (sqrt). The spearman’s correlation coefficients and P values for 97,853,055 pairs of genes commonly detected in both Ctrl and Exosc3 KO were calculated using the normalized matrices. The P values were corrected by Benjamini–Hochberg’s method. To robustly estimate the Spearman’s rank correlation coefficient and assess its statistical significance, we employed the permutation-based approach implemented in the perk R package,^131^ based on an appropriately studentized statistic. This method is particularly suitable for datasets with non-normal distributions.^152^ The permutation-based P value was determined by two sided tests and corrected by Benjamini–Hochberg’s method. Significant co-expression pairs were determined based on the criteria: Spearman correlation coefficient > 0.1, adjusted P value < 0.01, as well as permutation-based adjusted P value < 0.01. To analyze the correlations of pairs of SE genes (40,186 pairs), TE genes (6,791,455 pairs), and randomly selected genes, we calculated the fractions of significant co-expression pairs among SE and TE gene pairs and compared with the result of 1,000 times sampling of randomly selected pairs whose number was matched to SE gene pairs (Figure 7D). For correlation heatmaps, Genes were ordered by hierarchical clustering with Euclidean distance and Ward’s criterion (ward.D2). The Hi-C contact bias categories were determined by the ratio calculated as in Figure 6E (“Down”, Down ratio > 0.5; “Up”, Up ratio > 0.5) without filtering by compartment size. To visualize the expression correlation in scatter plots, we applied sqrt-transformed matrices to MAGIC.^130^ Since MAGIC cannot distinguish transcriptional burst from technical noise, and to avoid inducing spurious correlation due to over-smoothing, we used relatively compromised parameter settings; knn=2 and t=3.

## QUANTIFICATION AND STATISTICAL ANALYSIS

In Figures 1B, 1D (boxplots), 4C-D, 4G, 4I, 6D, 6G, S1D (box plots), S1E-F, S3F-G, S4E-G, S5B, and S6E-F, statistical significance was evaluated with Wilcoxon ranked sum test.

In Figures 1D (bar plots), 1I-J, S1D (bar plots), and S1G, two-sided Fisher’s exact test between Ctrl and Exosc3 KO conditions was performed to evaluate statistical significance (1D (bar plots), comparison between nuclei with or without large MED19 condensates; 1I, comparison between condensates with or without Pol II overlaps; 1J, comparison between nuclei with or without Pol II-overlapped large MED19 condensates; S1D (bar plots), comparison between nuclei with or without large Pol II condensates; S1G, comparison between nuclei with or without MED19-overlapped large Pol II condensates).

In Figures 2G-H, 2J-K, 3E-F, and S3D-E, statistical significance was evaluated using two-tailed *t*-tests with Welch’s correction. For S3D and S3E, we performed downsampling with a 0.2 sampling ratio thirty times, and the calibrated dwell times of the sample condition were pooled together for performing *t*-tests.

In Figures 2L-M, statistical significance was evaluated with chi-square test.

In Figures 4C-D and S4E-F, ChIP-seq signals were visualized in the following ranges: Region, scaled regions for SEs with ± 1 kb flanking regions; Peak, ± 3 kb regions centering on enhancer constituent summits. For details, see *Metaplot* in EXPERIMENTAL MODEL AND SUBJECT DETAILS.

In Figures 4E and 4H, PRO-seq signals or log2 fold-change (KO vs Ctrl) were visualized in the following ranges: 4E, ± 1kb regions centering on enhancer constituent summits; 4H, −2 kb to +10 kb regions surrounding the TSS of the genes. For details, see *Metaplot* in EXPERIMENTAL MODEL AND SUBJECT DETAILS.

In Figure 5G, Pearson’s correlation coefficient and BH-corrected P values (chi-square test) were calculated after binarizing the count matrices. Co-expressed eRNA-gene pairs were extracted using the following criteria: FDR < 0.05 and Pearson correlation coefficient > 0.1. For details, see *eRNA-gene co-expression analysis* in EXPERIMENTAL MODEL AND SUBJECT DETAILS.

In Figures 6F and S6G, statistical significance was evaluated with Kolmogorov–Smirnov test.

In Figures 7D-E, Spearman correlation coefficients, BH-corrected P values for the correlation, and BH-corrected permutation-based P values were calculated. Co-expressed gene pairs were extracted using the following criteria: Spearman’s r > 0.1, adjusted P value < 0.01, and adjusted P value from the permutation test < 0.01. For details, see *Gene-gene co-expression analysis* in EXPERIMENTAL MODEL AND SUBJECT DETAILS.

In all analyses, P values were corrected for multiple testing where appropriate.

## SUPPLEMENTAL TABLES

**Table S1, related to** ^16^**. sgRNA information.**

## SUPPLEMENTAL MOVIES

**Video S1. Simulation of intra-condensate exchange, related to Figure 2**.

**Video S2. Simultaneous observation of dSTORM movies of MED19 and single-molecules of Pol II in the living cells, related to Figure 3**.

**Video S3. Live-cell images showing co-bursting of endogenous Sox2 and Tet2 genes, related to Figure 7**

**Video S4. Live-cell images showing co-bursting of endogenous Nanog and Smarcad1 genes, related to Figure S7.**

## Supplementary Information

### Supplementary Figure Legends

**Figure S1.**
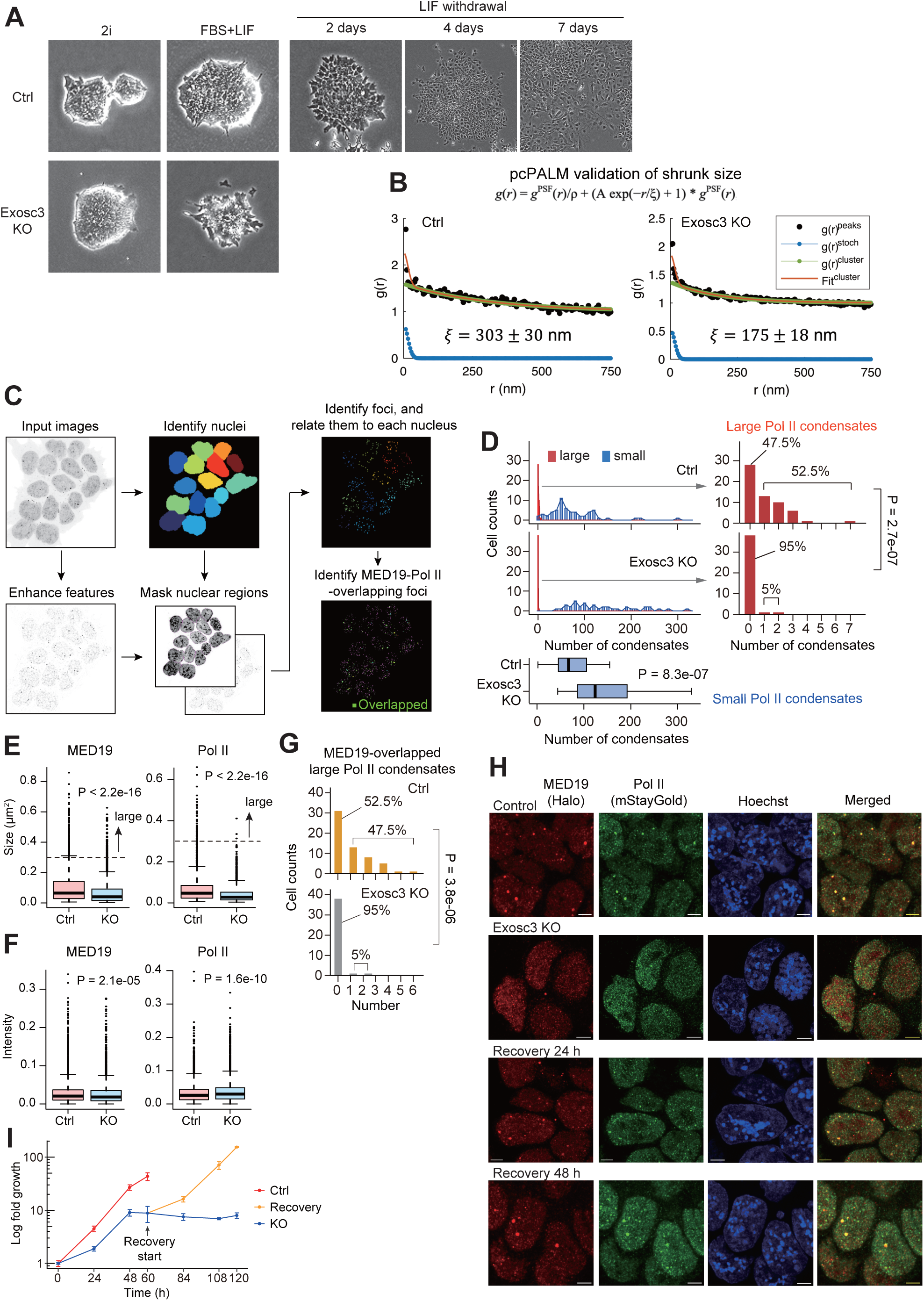
Impacts of RNA exosome depletion on the sizes and numbers of transcriptional condensates, related to Figure 1. (A) Bright-field images showing cell morphology under 2i and FBS+LIF conditions, before and after Exosc3 depletion, and before and after LIF withdrawal in FBS culture conditions in control cells. (B) A pair-correlation analysis of the spatial distribution from PALM images (pcPALM). The pair-correlation function as a function of radial distance *r* (nm) was computed. The plot shows the pair-correlation function calculated from individual peaks (black), and the curve fitted to a general function (orange), as well as the corrected correlation function for MED19 proteins (green) and the fluorophore stochastic contributions (blue). The characteristic cluster size ζ and the standard deviation are also shown. (C) Overview of the image analysis workflow using CellProfiler. (D) Histograms showing the number of small (< 0.3 µm^2^) and large (≥ 0.3 µm^2^) Pol II condensates per nucleus in control and Exosc3 KO cells. For small condensates, the x-axis is divided into bins of 5. Bottom boxplots and right bar plots represent distribution of small condensate counts per nucleus and enlarged view of large condensates, respectively. P value, Wilcoxon rank sum test (box plots) and Fisher’s exact test (bar plots). (E, F) Boxplots displaying size (E) and intensity (F) of MED19 and Pol II condensates. P value, Wilcoxon rank sum test. (G) Bar plots showing the number of large Pol II condensates co-partitioned with MED19. P value (bar plots), Fisher’s exact test comparing Ctrl vs KO (nuclei with or without large condensates). (H) Images showing the recovery of MED19 and Pol II condensates in the recovery phase of Exosc3 expression from the KO condition. Scale bar, 4 µm. (I) Growth curves of Exosc3 CKO/Halo-MED19/mSG-Pol II cells. Fold change of the number of viable cells were calculated against time 0. Mean ± sd (n = 3).

**Figure S2.**
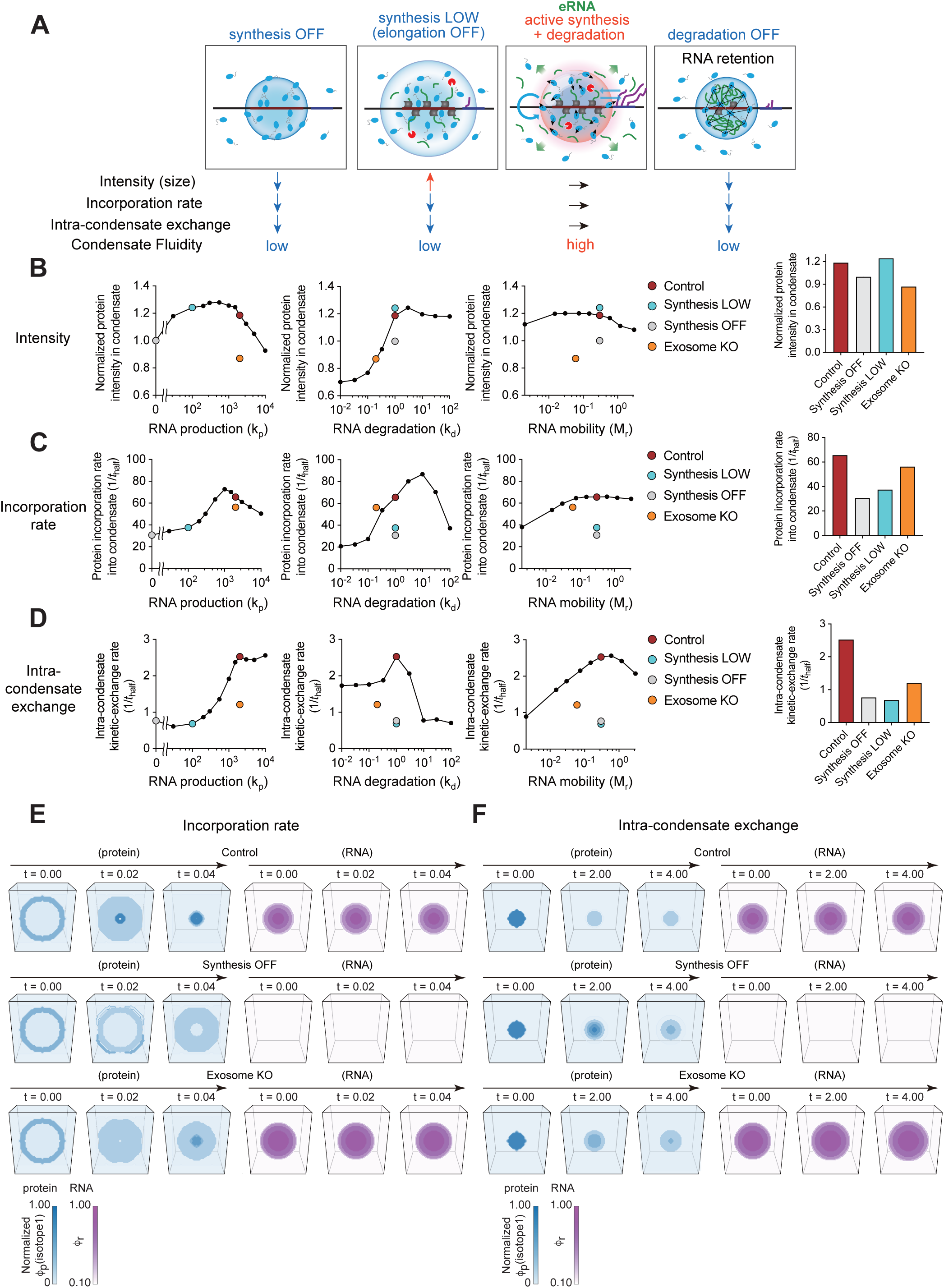
Simulations of alterations in fluidity and physical properties of transcriptional condensates in response to transcription and RNA exosome perturbations, related to Figure 2. (A) A schematic depicting the effects of inhibition of RNA transcription (synthesis OFF), inhibition of elongation (synthesis LOW), and inhibition of RNA destabilization on condensate fluidity. Summaries of the simulation results from Figures 2A-D and S2B-F are shown below. For details on the alterations in gene transcription, see Figure 4, STAR Methods, and the Discussion. (B-D) Simulation predictions of average intensity (B), incorporation rate (C), and intra-condensate exchange (D) of the condensate with varying RNA production rates (*kp*), RNA degradation rate (*kd*), and RNA mobility (*Mr*). Simulation results for control, inhibition of RNA transcription (synthesis OFF), inhibition of elongation (synthesis LOW), and Exosome KO condition are shown in both polyline graphs and bar plots. (E, F) Visualization of protein isotope 1 (blue) and RNA (magenta) concentration fields over simulation time for 3D simulations of incorporation rate (E) and intra-condensate exchange (F, see Video S1) in control, synthesis OFF, and Exosome KO conditions. Note that the overall protein field (isotope 1 + isotope 2) remains at the steady state (not shown).

**Figure S3.**
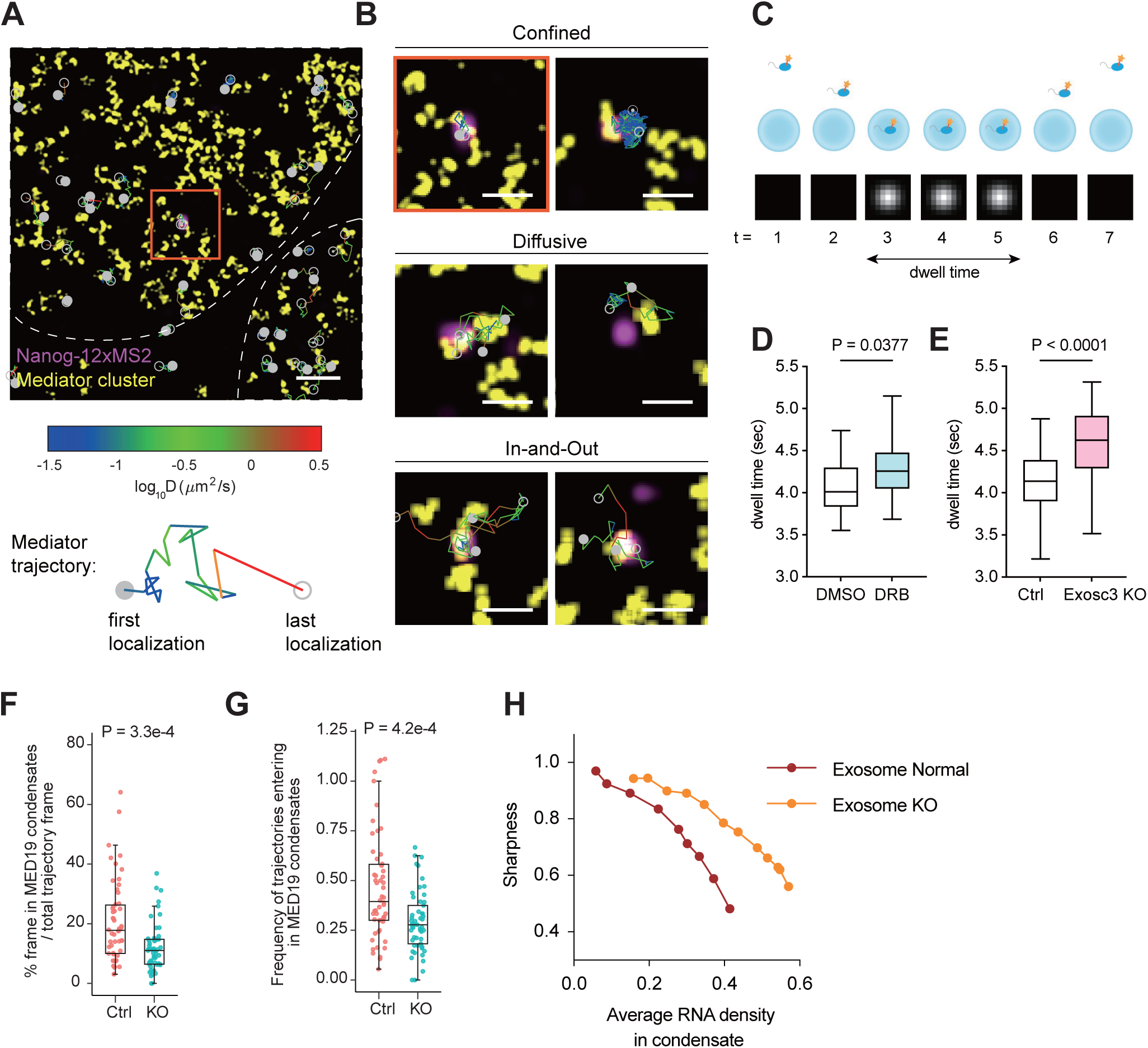
Stalled molecule exchange dynamics of transcriptional condensates in Exosc3 KO cells, related to Figures 2 **and 3**. (A) A representative image showing the trajectories of Mediator clusters around the *Nanog* transcription site. The colors of the trajectory lines indicate diffusion rates (Log D). (B) Representative diffusion patterns for confined, diffusive, and in-and-out trajectories around the *Nanog* transcription site. (C) A schematic depicting dwell time analysis of a Mediator molecule in condensates. (D, E) Boxplots illustrating changes in dwell time upon DRB treatment (20 min) (D) and Exosc3 depletion (E). P value, two-tailed *t*-test. (F) Results of single-molecule tracking of Pol II in the combination of dSTORM super-resolution imaging of MED19 (see Video S2). Boxplots show the percentage of frames where Pol II trajectories are detected in MED19 condensates relative to the total trajectory frames. Each dots represent the cells. P value, Wilcoxon rank sum test. (G) Boxplots showing the frequency of Pol II trajectories entering in MED19 condensates. P value, Wilcoxon rank sum test. (H) The relationship between average RNA density in condensates and condensate sharpness under normal and Exosome KO conditions in phase-field simulations.

**Figure S4.**
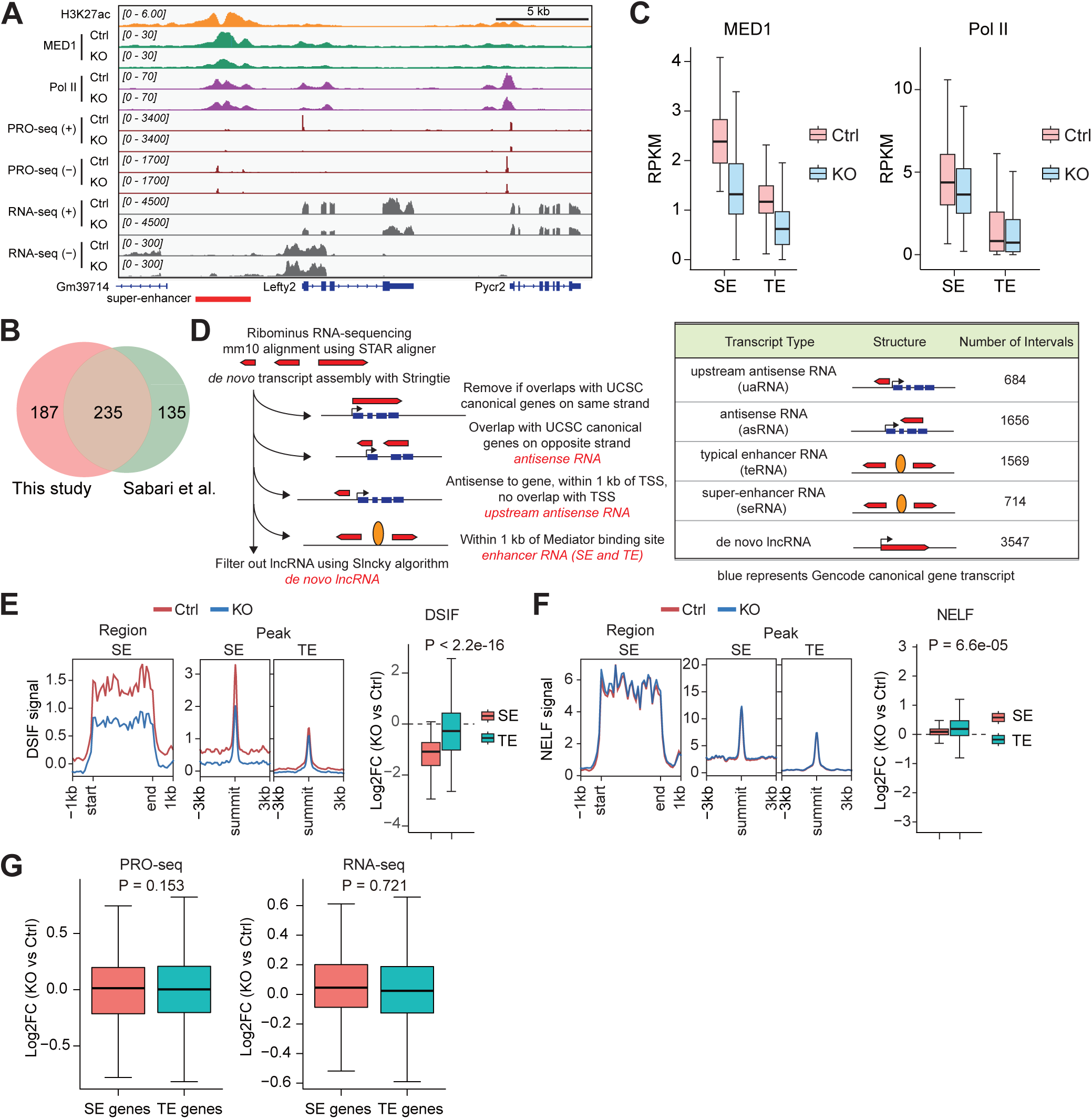
Multi-omics analysis in Exosc3 KO cells, related to Figure 4. (A) Additional representative IGV snapshots showing ChIP-seq, PRO-seq, and RNA-seq signals of Ctrl and Exosc3 KO cells around SE and the SE-nearest gene (*Lefty2*). (B) Venn diagram showing the substantial overlap of SEs identified in our study and a previous study (*Sabri et al.*, 2018),^23^ both of which were performed in 2i conditions. (C) Boxplots displaying the distribution of the input-subtracted MED1 (left) and Pol II (right) occupancies within SE and TE regions in control and Exosc3 KO cells. (D) Strategy for *de novo* transcriptome assembly and the resulting number of intervals. The image is adapted from our previous study^43^ and modified. (E, F) Metaplots showing mean input-subtracted DSIF (E) and NELF (F) ChIP-seq signal intensities across SE regions and a 6 kb window centered on SE and TE constituents (left). Boxplots represent the distribution of the log2-fold change (Exosc3 KO vs Ctrl) in ChIP-seq signals within SE and TE regions (right). P value, Wilcoxon rank sum test. (G) Boxplots displaying the log2-fold PRO-seq and RNA-seq signal changes (Exosc3 KO vs Ctrl) in SE and TE genes. P value, Wilcoxon rank sum test.

**Figure S5.**
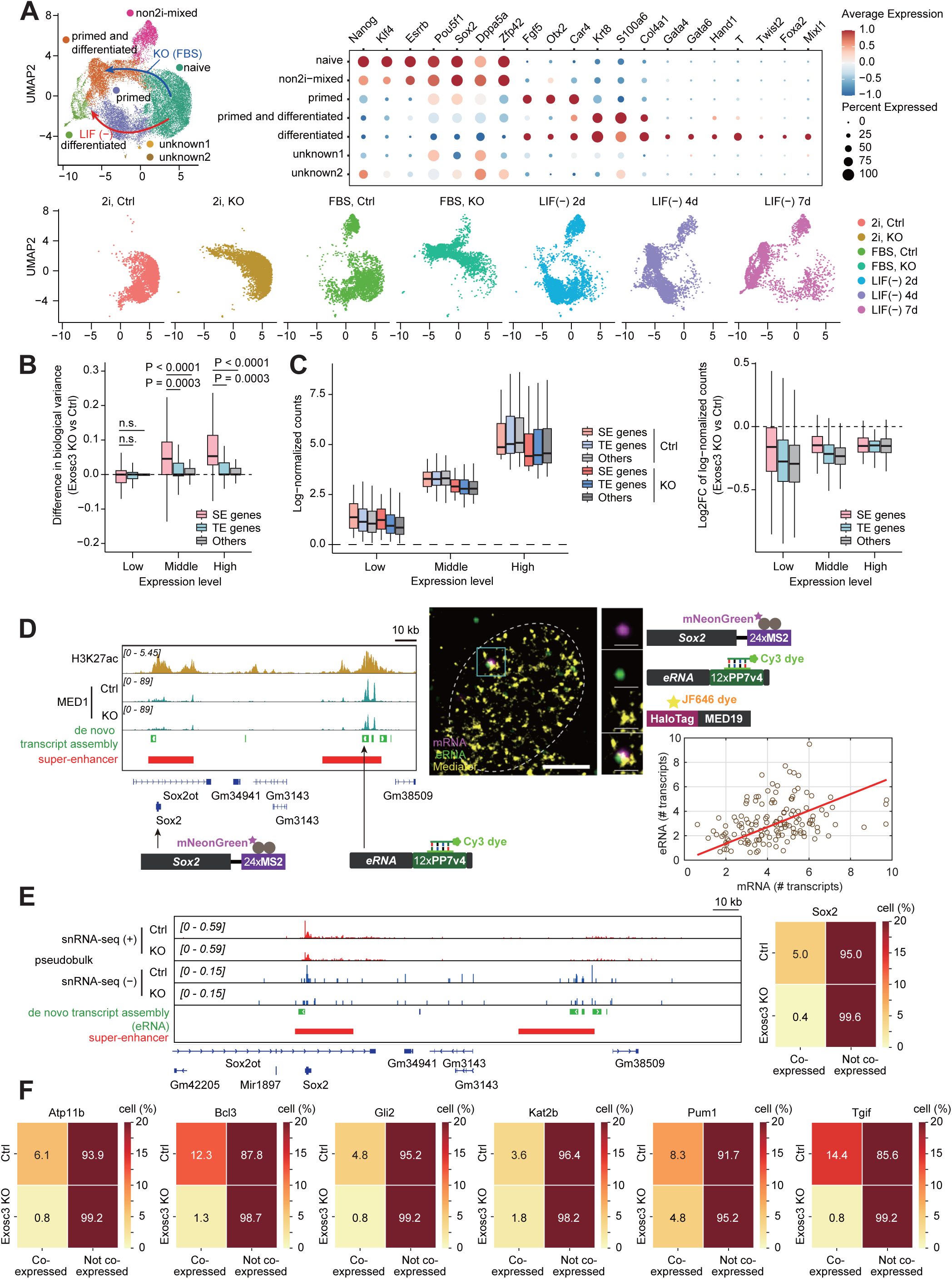
Pluripotency gene expression is maintained in Exosc3 KO cells, while eRNA-gene co-expression is lost for SE genes, related to Figure 5. (A) Detailed data of 3′-scRNA-seq analysis demonstrating cell classification and differentiation status, related to Figure 5B. Dot plots illustrate the expression of representative pluripotency and differentiation marker genes. UMAP data from Figure 5B is colored and split by the culturing conditions (bottom). (B) Boxplots showing differences in biological variance in the expression of SE-nearest, and TE-nearest, and other genes in 3′-scRNA-seq analysis, according to average expression level. P value, Wilcoxon rank sum test. (C) Boxplots showing the log-normalized counts of SE-nearest, and TE-nearest, and other genes in 3′-scRNA-seq analysis (left) and the log2-fold change (Exosc3 KO vs Ctrl) of the genes (right), according to average expression level. (D) IGV snapshots showing the *Sox2* gene and its SE eRNA fused with MS2 and PP7 repeats (left), additional representative super-resolution image showing *Sox2* gene and eRNA co-bursting around MED19 clusters (middle), and a scatter plot showing correlation between the number of Sox2 mRNAs and eRNAs (right). Scale bar, 3 µm; cropped and zoomed images, 1 µm^2^ area centering the bursting site. (E) Additional representative IGV snapshots showing pseudobulk 5′-snRNA-seq signals of control and Exosc3 KO cells around SE and the SE-nearest gene (*Sox2*). The right heatmap illustrates the proportion of gene-expressing cells that either co-expressed eRNAs (Co-expressed) or did not (Not co-expressed). (F) Heatmaps showing additional examples (Atp11b, Bcl3, Gli2, Kat2b, Pum1, and Tgif) of the proportion of SE-nearest gene-expressing cells that either co-expressed eRNAs or did not.

**Figure S6.**
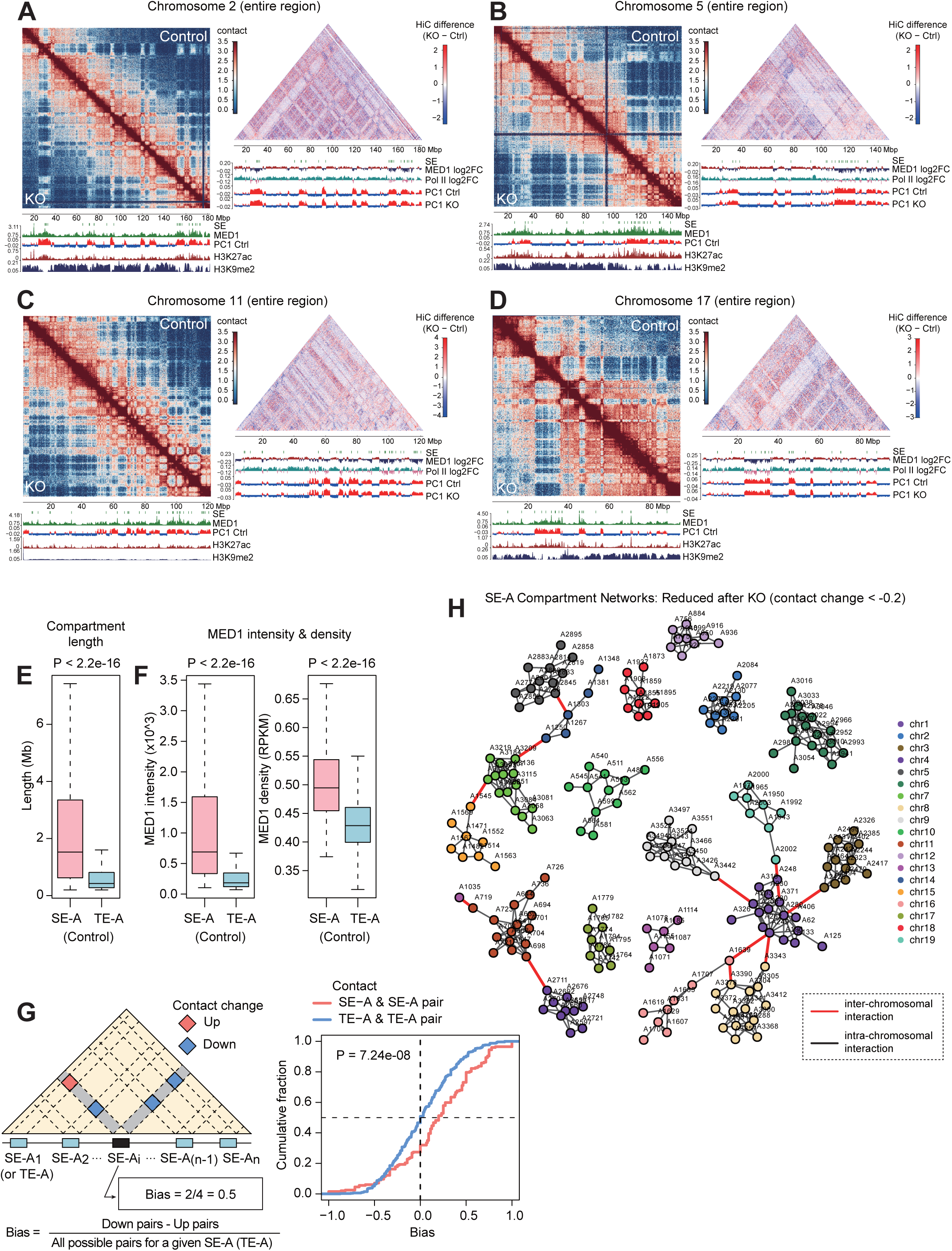
Altered Hi-C interactions in Exosc3 KO cells, related to Figure 6. (A-D) Contact matrices entire chromosomes 2 (A), 5 (B), 11 (C), and 17 (D) in control and Exosc3 KO conditions (left) and the differential contact matrices (right) showing the changes in the interaction after Exosc3 KO at 200 kb resolution. Bottom tracks show ChIP-seq signals for MED1 (Ctrl), H3K27ac, and H3K9me3 (left), the log2 FC of MED1 and Pol II ChIP-seq signals (Exosc3 KO vs Ctrl), PC1 scores, and the position of SEs (right), respectively. H3K27ac and H3K9me3 ChIP-seq data in 2i-cultured mESC are from *Yang et al*., 2019.^105^ (E) Boxplots showing the length of SE-A and TE-A compartments in control cells. P value, Wilcoxon rank sum test. (F) Boxplots showing MED1 ChIP-seq signal intensity and density for SE-A and TE-A compartments in control cells. P value, Wilcoxon rank sum test. (G) A diagram shows contact changes between SE-As (or TE-As) and how the bias of “Down” vs “Up” was calculated for a given region (left). CDF plots of the bias for SE-A/SE-A and TE-A/TE-A pairs are shown (right). P value, Kolmogorov-Smirnov test. (H) SE-A interaction network in the autosomal chromosomes, whose contacts are sensitive to Exosc3 KO. Gray and red edges indicate intra-chromosomal and inter-chromosomal SE-A connections, respectively.

**Figure S7.**
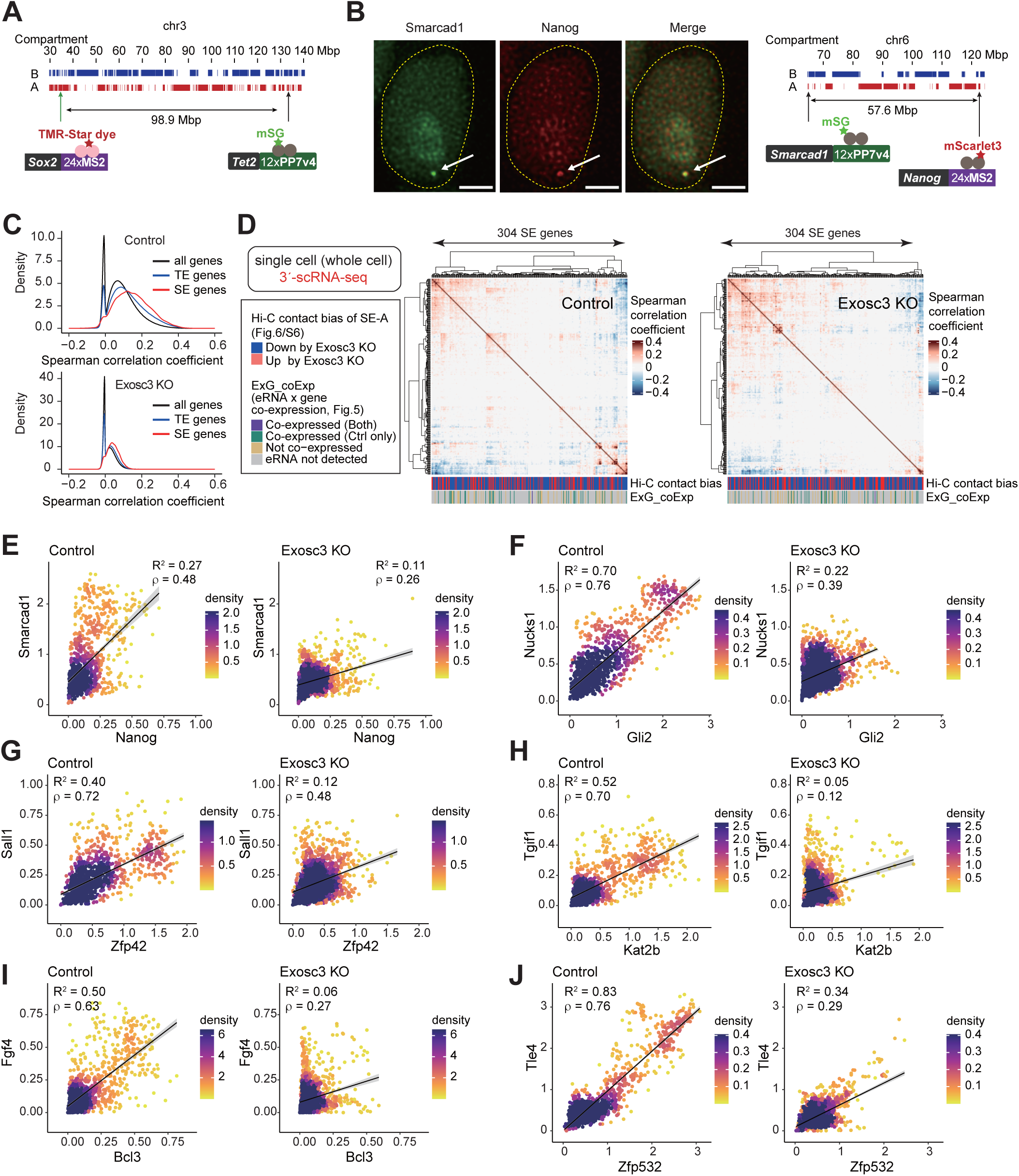
5′-snRNA-seq outperforms 3′-scRNA-seq in gene-gene correlation analysis as a proxy for transcriptional activity, related to Figure 7. (A) A schematic showing the genomic organization of *Sox2* and *Tet2* gene loci. (B) Representative live-cell images showing co-bursting of endogenous *Nanog* and *Smarcad1* genes in a control cell (left, see Video S4), and a schematic showing the genomic organization of *Nanog* and *Smarcad1* gene loci (right). Arrows indicate the co-bursting. We observed substantial temporal overlap between Nanog and Smarcad1 bursts (Nanog: 44 and 56-72 min, Smarcad1: 28-76 min). Scale bars, 2 µm. (C) Density plots depicting the distribution of Spearman’s correlation coefficients for SE gene, TE gene, and all possible gene pairs in Ctrl (top) and Exosc3 KO (bottom) cells. (D) Heatmaps showing Spearman’s correlation coefficients among SE genes in Ctrl and Exosc3 KO cells, calculated using 3′-scRNA-seq data. Genes are ordered by hierarchical clustering with Euclidean distance and Ward’s criterion (ward.D2). The annotations of the columns are as in Figure 7F: The Hi-C contact bias categories were determined by the ratio calculated as in Figures 6E-F (“Down”, Down ratio > 0.5; “Up”, Up ratio > 0.5). ExG_coExp represents the eRNA-gene co-expression status determined in Figures 5F-G. (E-J) Examples of co-transcribed SE gene pairs; Nanog-Smarcad1 (E), Gli2-Nucks1 (F), Zfp42-Sall1 (G), Kat2b-Tgif1 (H), Bcl3-Fgf4 (I), and Zfp532-Tle4 (J). Sqrt-normalized counts from 5′-snRNA-seq data were processed using the MAGIC algorism with modest smoothing parameters (knn = 2, t = 3). The resulting values for the indicated gene pairs in Ctrl and Exosc3 KO were plotted. Lines represent linear regression with 95% confidence intervals (shaded areas). *R*^2^, coefficient of determination; ρ represents Spearman’s correlation coefficient.

**Video S1. Simulation of intra-condensate exchange rate, related to Figure 2**. Visualization of protein isotope 1 (blue) concentration fields over simulation time for 3D simulations of intra-condensate exchange rate in control, synthesis OFF, and Exosome KO conditions are shown.

Binarized domains of MED19 clusters (red) and single-molecule tracking data of the Pol II (green) are shown.

24×MS2 (3′ UTR) and 12×PP7v4 were fused to Sox2 and Tet2 genes, respectively, in mESCs expressing MCP-SNAP and PCP-mSG. White circle indicates the position where burst occurs. Scale bar, 2 µm.

**Video S4. Live-cell images showing co-bursting of endogenous Nanog and Smarcad1 genes, related to Figure 7**.

24×MS2 (3′ UTR) and 12×PP7v4 were fused to Nanog and Smarcad1 genes, respectively, in mESCs expressing MCP-mScarlet3 and PCP-mSG. White circle indicates the start of the burst. Scale bar, 2 µm.

